# Targeted and efficient AAV therapy for neuroblastoma via direct capsid-antibody coupling

**DOI:** 10.1101/2025.11.12.687993

**Authors:** Mirko Luoni, Serena G. Giannelli, Stefano Parracino, Francesca Morosi, Elisa Ventura, Agostina Di Pizio, Sharon Muggeo, Angelo Iannielli, Tamara Canu, Tommaso Russo, Antonio Esposito, Fabio Pastorino, Vania Broccoli

## Abstract

Adeno-associated viral (AAV) vectors are widely used in gene therapy for their versatility and safety, but their broad tropism limits cell-specific applications such as targeting primary or metastatic tumor cells. To address this, we developed AAV-STITCH, a strategy using SpyTag technology to covalently attach polypeptides to the AAV capsid. This allows precise, dose-dependent coupling of an anti-GD2 ScFv to a galactose-binding-deficient AAV9-W503A capsid, redirecting tropism specifically to GD2-expressing neuroblastoma (NB) cells. In pseudometastatic xenograft mouse models, AAV-STITCHαGD2 selectively transduced NB tumor cells without transduction of healthy tissues. Furthermore, delivery of a suicide gene via AAV-STITCHαGD2 suppressed tumor growth and extended survival in mice with subcutaneous and pseudometastatic NB xenografts. These findings establish the feasibility of engineering AAVs with cell-type-specific transduction properties, providing a powerful and adaptable platform for the selective elimination of NB tumor cells. This technology marks a meaningful advance toward next-generation, targeted cancer therapies with strong clinical potential.

## Introduction

Neuroblastoma (NB) is the most frequent and aggressive pediatric extracranial solid tumor, responsible for 15% of childhood cancer deaths (1). Approximately half of all NB patients are classified with high-risk (HR) disease typical with distant site metastases at diagnosis. Despite recent multimodal therapeutic regimens, including chemotherapy, radiation, surgical resection and autologous stem cell transplant, the 5-year survival still remains below 50% in the high-risk group (2). Moreover, children in whom first-line therapy fails have a very low probability of recovery with subsequent treatments, and they have a dismal prognosis with a long-term survival below 10%. Thus, more effective therapies are urgently needed to offer a better prognosis to these patients. The discovery that the disialoganglioside GD2 is highly expressed in NB and other cancers of neuroectodermal origin, while being rarely present in healthy peripheral tissues has spearheaded the development of immunotherapies for these malignancies. The introduction of GD2 monoclonal antibodies, has provided a new therapeutic option, improving the survival rate in high-risk NB patients (3). However, serious adverse events, including acute pain, have limited the development of this therapy (4). GD2 was one of the first tumor-associated antigens targeted by CAR T cells in humans, and clinical trials support the efficacy of GD2-redirected CAR T cells with encouraging clinical activity in first patients (5-7). However, high cellular doses can lead to systemic inflammation posing important obstacles to be yet overcome. Moreover, some GD2 expression is detectable on neuronal cell bodies in different brain regions (8), raising concerns for neurotoxicity, although brain-related adverse effects have not yet observed in the treated patients. More in general, high-risk NBs present limited T-cell infiltration, MHC-I downregulation and low mutational burden leading to a significant immunosuppressive tumor microenvironment which can challenge the outcome of CAR T-cell therapies (2). Thus, despite these important recent therapeutic advances that have improved the prognosis of high-risk NB patients, it remains urgent the need for additional and synergic medical products.

AAVs are non-pathogenic viruses that can infect both dividing and quiescent cells for long-term gene expression (9). Recombinant AAVs are produced at high yield in large-scale manufacturing and have broad tropism making them the most widely used vectors for gene therapy (10). In fact, numerous gene therapy clinical trials worldwide are currently in development using AAVs as delivery system for a vast number of human diseases. However, AAV gene therapy applications in cancer are scarce given the general non-specific tropism of these viruses. In fact, AAV native serotypes show broad infectivity in multiple tissues, although with some preferred cell-type tropism. Despite the success in modulating the AAV infectivity pattern by targeted capsid engineering, it is missing a convenient approach to attain cell-type specific viral transduction (11). We developed a new and powerful strategy to covalently couple antibody fragments on the AAV capsid surface in order to control viral infectivity for achieving selective transduction of NB cancer cells. These modified capsids demonstrated effective and safe anti-tumoral activity in xenograft NB mouse models, validating their strong translational therapeutic potential.

## Results

### Efficient and direct coupling of an αGD2-scFv to the AAV9 capsid by the SpyTag technology

To establish an effective and modular approach for coupling antibody fragments to AAV capsids, we leveraged SpyTag technology. This system relies on two small reactive protein fragments, SpyTag and SpyCatcher, which spontaneously form a covalent isopeptide bond with high affinity (12). Given that the SpyTag fragment consists of only 13 amino acids, we hypothesized that its integration into AAV variable regions would not destabilize the AAV structure. Thus, the SpyTag sequence, flanked by spacer elements, was cloned after amino acid 453 (the VP1 position) within the variable region IV of the AAV9 capsid, which has been shown to be an editable site in the capsid without altering the virus productivity and infectivity (13) (Fig. 1a). We selected the AAV9 capsid serotype for its high productivity yield. Moreover, we recently showed that the AAV-W503A mutant capsid, previously described to abolish its binding to galactose, is fully infection deficient (14). Thus, we reasoned that this AAV9 mutant capsid represents a perfect neutral packaging vector, whose tropism will be solely dictated by the peptides grafted onto its surface. Next, we fused the SpyCatcher with the GD2 scFv antibody fragment (14g2a) (a kind gift of M. Brenner) with an optimized linker sequence (5) (Fig. 1a). AAV production was carried out with standard transfection of HEK293 cells using AAV-producing plasmids and the SpyCatcher-αGD2 vector, we termed this strategy AAV-STITCH for SpyTag Targeted Coupling. When the virus and SpyCatcher-αGD2 were produced separately and subsequently incubated together, a significant fraction of unbound SpyCatcher-αGD2 remained in the sample (post-production protocol) (Suppl. Fig. 1a). In contrast, co-producing the virus and the SpyCatcher-αGD2 fusion in the same cells, followed by purification on iodixanol gradient, resulted in the efficient removal of unbound SpyCatcher-αGD2, which was largely excluded from the final viral preparation (co-expression protocol) (Suppl. Fig. 1b). Therefore, we chose the latter procedure to avoid potential interference by the contaminant free SpyCatcher-αGD2 molecules in the AAV preparation.

**Figure 1.**
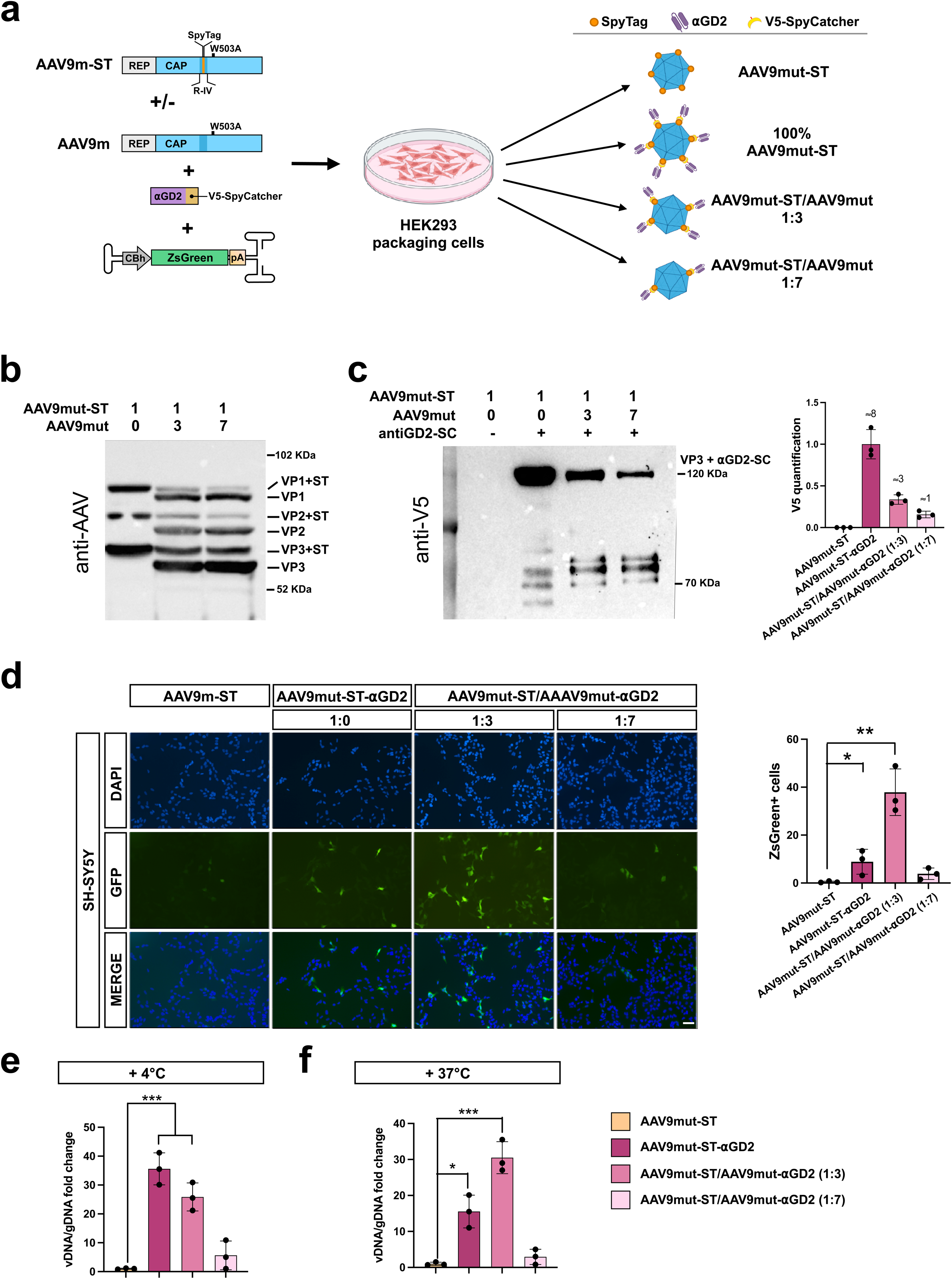
Production and functional validation of mosaic AAV-STITCH vectors. **a.** Schematic of the experimental protocol used for AAV-STITCH and mosaic capsid production by co-expressing plasmids encoding the AAV components together with the SpyCatcher fused to the anti-GD2 scFv (⍺GD2-SC). **b**. Western blot with anti-AAV antibody showing the capsid composition of SpyTag-modified and mosaic AAV preparations. **c**. On the right, western blot using an anti-V5 antibody to assess the coupling rate of the ⍺GD2-SC fusion to non-mosaic and mosaic (1:3 and 1:7; AAV9mut-ST/AAV9mut) AAV-STITCH capsids. On the left, densitometric quantification of the bands and calculation of the average number of ⍺GD2-SC molecules conjugated per viral capsid. n=3 independent biological replicates**. d**. On the left, immunofluorescence using anti-ZsGreen antibody on SH-SY5Y cells transduced with control AAV9mut-ST and ⍺GD2-SC (non-mosaic 1:0 and mosaic 1:3 - 1:7). Scale bar: 100 µm. On the right, bar graph showing the quantification of ZsGreen+ cells over total cells in the different experimental conditions. N = 3 independent biological replicates. **e.** RT-qPCR quantification of cell membrane–bound viral genomes (4°C assay) following incubation with control AAV9mut-ST alone or conjugated with ⍺GD2-SC (non-mosaic 1:0 and mosaic 1:3 - 1:7). n = 3 independent biological replicates. **f**. RT-qPCR quantification of viral genome copies in SH-SY5Y cells transduced with control AAV9mut-ST alone or conjugated with ⍺GD2-SC (non-mosaic 1:0 and mosaic 1:3 - 1:7) at 37°C. n = 3 independent biological replicates. Error bars represent SD. *p<0.05; **p<0.01; ***p < 0.001, One-way ANOVA with Tukey’s multiple comparisons correction.

We opted to produce self-complementary AAVs since they assure a faster and higher transgene expression which is crucial when targeting rapidly dividing cancer cells (15). Multiple independent virus productions revealed that the viral titer remained consistent, indicating that the co-expression protocol does not compromise virus yield (Suppl. Fig. 1c). To investigate the incorporation of the SpyCatcher-αGD2 into the capsid, we performed Western blot analysis using an antibody specific for AAV capsid proteins. We observed that SpyCatcher-αGD2 predominantly bound to the VP3 protein (Suppl. Fig. 1d). Densitometric quantification of the Western blot signals revealed that approximately 13 ± 3% of VP3 subunits were conjugated to SpyCatcher-αGD2. Given that each AAV capsid contains ∼50 VP3 molecules, we estimated that an average of 7-8 SpyCatcher-αGD2 molecules were conjugated per capsid (Suppl. Fig. 1d). When testing transduction efficiency in GD2+ SHSY-5Y neuroblastoma cells, we found that the unconjugated AAV was unable to transduce the cells (Suppl. Fig. 1e). In contrast, the antibody-conjugated AAV was able to mediate transduction, albeit with an efficiency below 10%. (Suppl. Fig. 1e). We hypothesized that the high density of SpyCatcher-αGD2 molecules on the viral surface might create steric hindrance, interfering with the interaction between the AAV capsid and essential co-receptors required for cellular entry. To test this hypothesis, we applied the same production protocol to generate mosaic capsids by mixing Rep-Cap plasmids containing or lacking the SpyTag sequence at different ratios (1:3 and 1:7) (Fig. 1a,b). This approach effectively reduced the number of SpyCatcher-αGD2 molecules conjugated to VP3 subunits from approximately 7–8 in the non-mosaic condition to 2–3 in the 1:3 ratio, and to fewer than 1 in the 1:7 ratio, based on V5 Western blot quantification (Fig. 1c). To determine these values, we first estimated the number of SpyCatcher-αGD2 molecules in the non-mosaic condition (set as ∼7 copies) (Suppl. Fig. 1d) and then normalized the band intensities in mosaic conditions accordingly (Fig. 1c). We then tested the transduction efficiency of the mosaic capsids *in vitro* and found that the 1:3 condition resulted in more than a 3-fold increase in transduction compared to the non-mosaic capsids (Fig. 1d). To assess viral binding independently of internalization, we performed a binding assay at 4°C, a condition under which viral entry is blocked, and observed that mosaic capsids exhibited reduced binding to the membrane of SH-SY5Y cells (Fig.1e). However, when analyzing viral vector copy number in cells following transduction at 37°C, we found that the 1:3 mosaic capsids resulted in a higher number of viral genomes per cell, consistent with the increased expression of the ZsGreen reporter observed by immunofluorescence (Fig. 1f). Based on these findings, we selected the 1:3 mosaic capsid configuration and designated it as AAV-STITCHαGD2. We then expanded its characterization to further evaluate its specificity and transduction efficiency across different neuroblastoma (NB) cell lines. Notably, the AAV-STITCHaGD2 efficiently transduced GD2+ NB cancer cell lines, such as SH-SY5Y, LAN-5, IMR-42 and HTLA-230, as shown by expression of the ZsGreen viral transgene (Suppl. Fig. 2). Conversely, GI-ME-N and SK-N-SH NB tumor cells, which are negative for GD2, resulted resistant to AAV-STITCHaGD2 transduction (Suppl. Fig. 2). We assumed that beyond its therapeutic potential, the AAV-STITCH system can also serve as a powerful tool to map the spatial distribution of antigens that are difficult to detect by conventional immunohistochemistry, such as GD2. In fact, despite its clinical relevance, GD2 expression remains poorly characterized, with only limited evidence on its presence in select cell types within the adult brain (16,17). To address this gap, we leveraged the ZsGreen-expressing AAV-STITCHαGD2 system to trace GD2 distribution in mouse brain organotypic slice cultures, as the mutant AAV9 vector is unable to cross the blood–brain barrier (Suppl. Fig. 3a). Viral transduction was observed in discrete neural cell types, including the granular and Purkinje cells of the cerebellum, as well as oligodendrocytes in the corpus callosum, cell populations previously suggested to express GD2 (16,17) (Suppl. Fig. 3b). These findings provide the first evidence that the αGD2 moiety is sufficient to redirect viral tropism, enabling specific transduction of both healthy and malignant GD2⁺ neural cells.

### AAV-STITCHαGD2 show selective infectivity of neuroblastoma tumor cells *in vivo*

To stringently assess any residual transduction of the AAV-STITCHαGD2 in healthy tissues, we inoculated a viral dose of 5x10e10 vg in adult wild-type C57BL/6 mice and analyzed the expression of the viral gene in many tissues normally infected by the native AAV9 serotype. Notably, ZsGreen fluorescence was undetectable in all tissues that normally are efficiently transduced by AAV9 including liver, heart and muscles (Fig. 2). Failed transduction was also observed with the AAV9m-ST capsid, indicating that αGD2 coupling did not mitigate the impaired infectivity resulting from the targeted AAV-W503A mutation (Fig. 2). The only notable exception was the dorsal root ganglia (DRG) neurons, which were efficiently transduced by AAV-STITCHαGD2, a finding consistent with previous reports describing high GD2 expression levels in this neuronal population (16,17) (Fig. 2).

**Figure 2.**
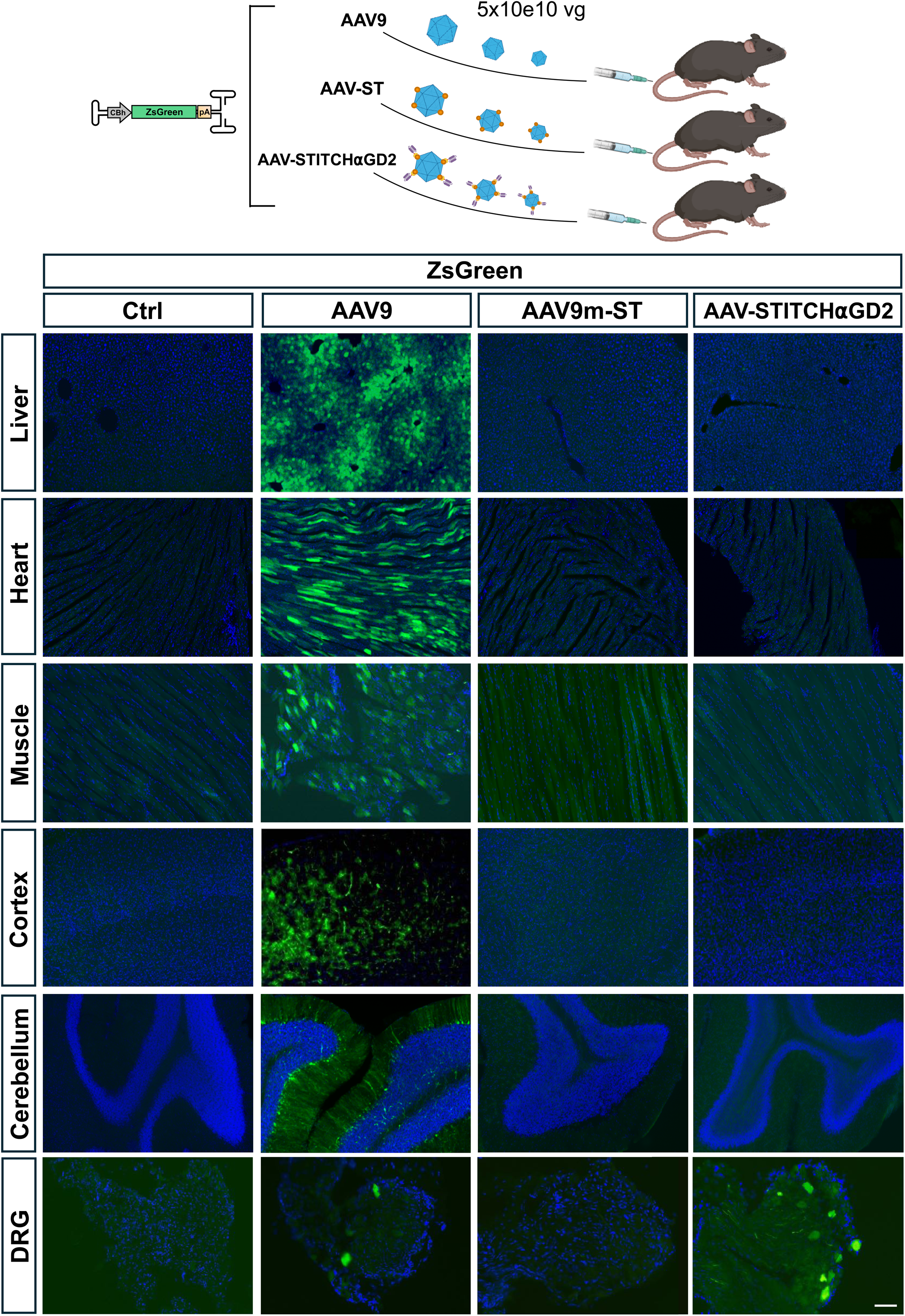
Biodistribution of AAV-STITCH⍺GD2 following systemic delivery in mice. **Top:** schematic of the experimental protocol. Briefly, we assessed the biodistribution of AAV-STITCH⍺GD2 carrying the ZsGreen transgene following systemic injection in C57BL/6 mice, compared to control injections with mutated AAV9-ST. **Bottom**: Representative immunofluorescence images showing ZsGreen expression, detected with an anti-ZsGreen antibody, in various organs to evaluate AAV-mediated transduction. Scale Bar: 100 µm.

Next, we established a pseudometastatic NB xenograft mouse model, by intravascular inoculation of GD2+ SH-SY5Y tumor cells in NSG mice (Fig. 3a). This model rapidly developed neuroblastoma metastases, with a notable enrichment in the liver within a few weeks (18). To evaluate the specific targeting efficiency of the AAV-STITCHαGD2 in neuroblastoma tumor cells, NSG mice were administered with this vector or a control virus (AAV9m-ST) two weeks following the inoculation of SH-SY5Y NB cells. Four days later, livers were then isolated and analyzed for the viral fluorescence signal. As shown in Fig. 3b, metastatic tissue was strongly positive to viral ZsGreen fluorescence. Notably, single NB tumor cells were successfully targeted by AAV-STITCHαGD2, while the surrounding healthy hepatic parenchyma remained completely untransduced (Fig. 3c). Conversely, no evidence of transduction of either malignant or healthy liver tissue was recorded using the control AAV9m-ST virus (Fig. 3b,c). Thus, AAV-STITCHαGD2 acquired a very specific tropism for NB tumor cells sparing the infection of healthy tissue in the liver.

**Figure 3.**
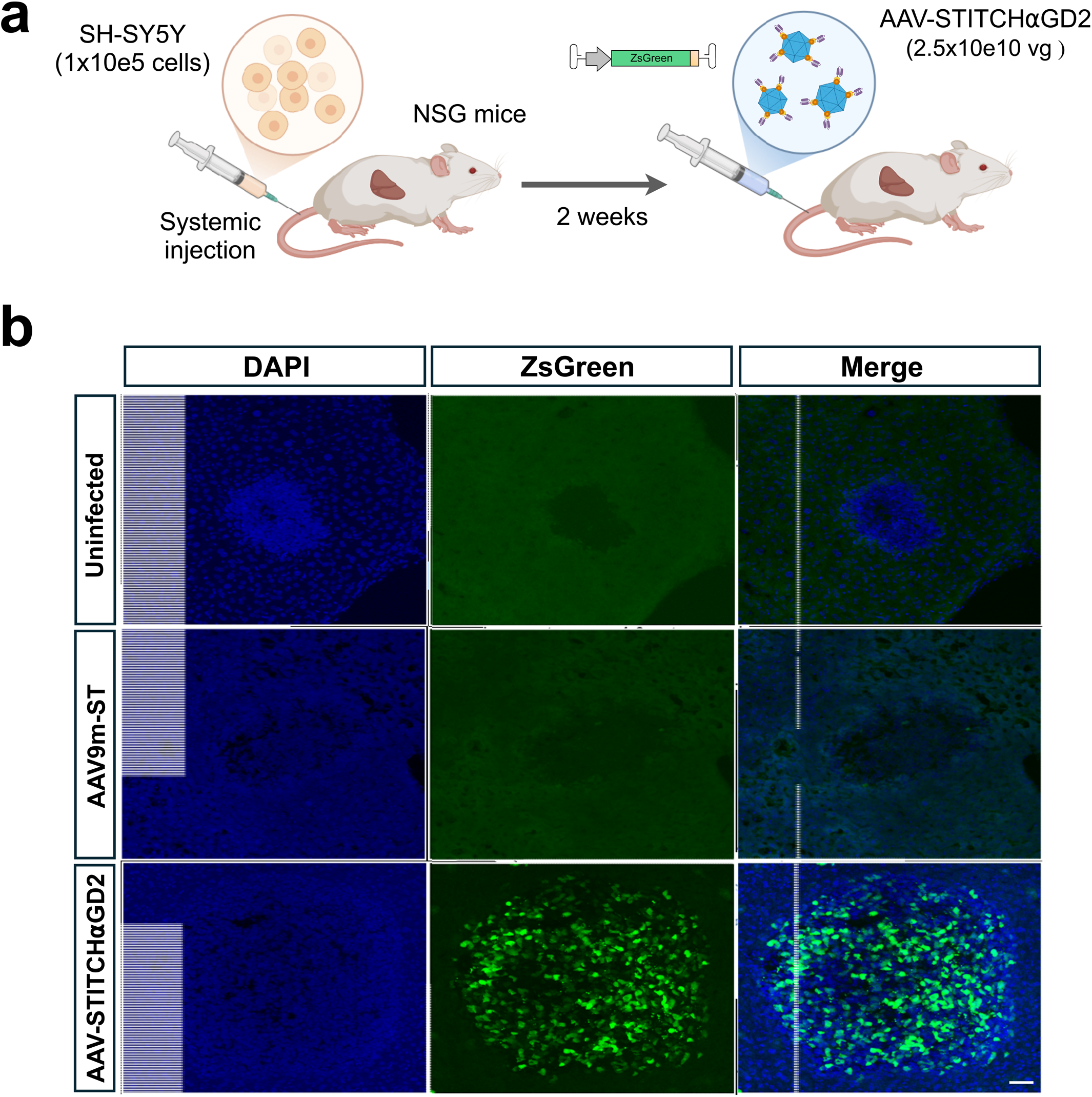
AAV-STITCHαGD2 targets liver metastasis in pseudometastatic neuroblastoma model. **a**. Scheme of the experimental protocol. In brief, 1x10e5 SH-SY5Y cells were systemically injected into adult NGS mice (pseudometastatic model). Two weeks later, AAV-STITCHαGD2 or AAV-ST as control carrying Zsgreen transgene were inoculated in the tail vein using 2.5x10e10 vg/mouse. Finally, treated mice were sacrificed three days after viral injection. **b**. Representative immunofluorescence for anti-ZsGreen in livers of pseudometastatic mice untreated or treated with AV-STITCHαGD2 or AAV-ST expressing Zsgreen transgene. Scale Bar: 100 µm.

### TK-expressing AAV-STITCHαGD2 has strong anti-tumoral effects

We then equipped the AAV-STITCHaGD2 with the Herpes Simplex Virus thymidine kinase (TK) gene which converts ganciclovir (GCV) into a toxic product, inducing irreversible apoptotic cell death as confirmed on *in vitro* assays with SH-SY5Y cells (Suppl. Fig. 4).

To enable convenient monitoring of AAV-STITCHαGD2 effects, we generated SH-SY5Y cells stably expressing Luc2, allowing longitudinal intravital bioluminescence imaging of subcutaneous xenografts (Fig. 4a). Next, we transplanted 1x10e6 Luc2-SH-SY5Y cells into the flank of NSG mice and two weeks later, animals were selected for comparable luciferase signal and divided into two groups which received a viral dose of 1x10e11 vg of either TK-AAV-STITCHaGD2 or the control vector AAV9m-ST (Fig. 4a). Notably, AAV-STITCHαGD2-TK robustly suppressed the growth of subcutaneous xenografts compared to the mock condition, as demonstrated by bioluminescence signal measurements over the 7-day observation period (Fig. 4b,c). Autoptic histological analysis confirmed that NB subcutaneous grafts treated with AAV-STITCHαGD2-TK were markedly reduced in size compared to tissues exposed to the control virus (Fig. 4d).

**Figure 4.**
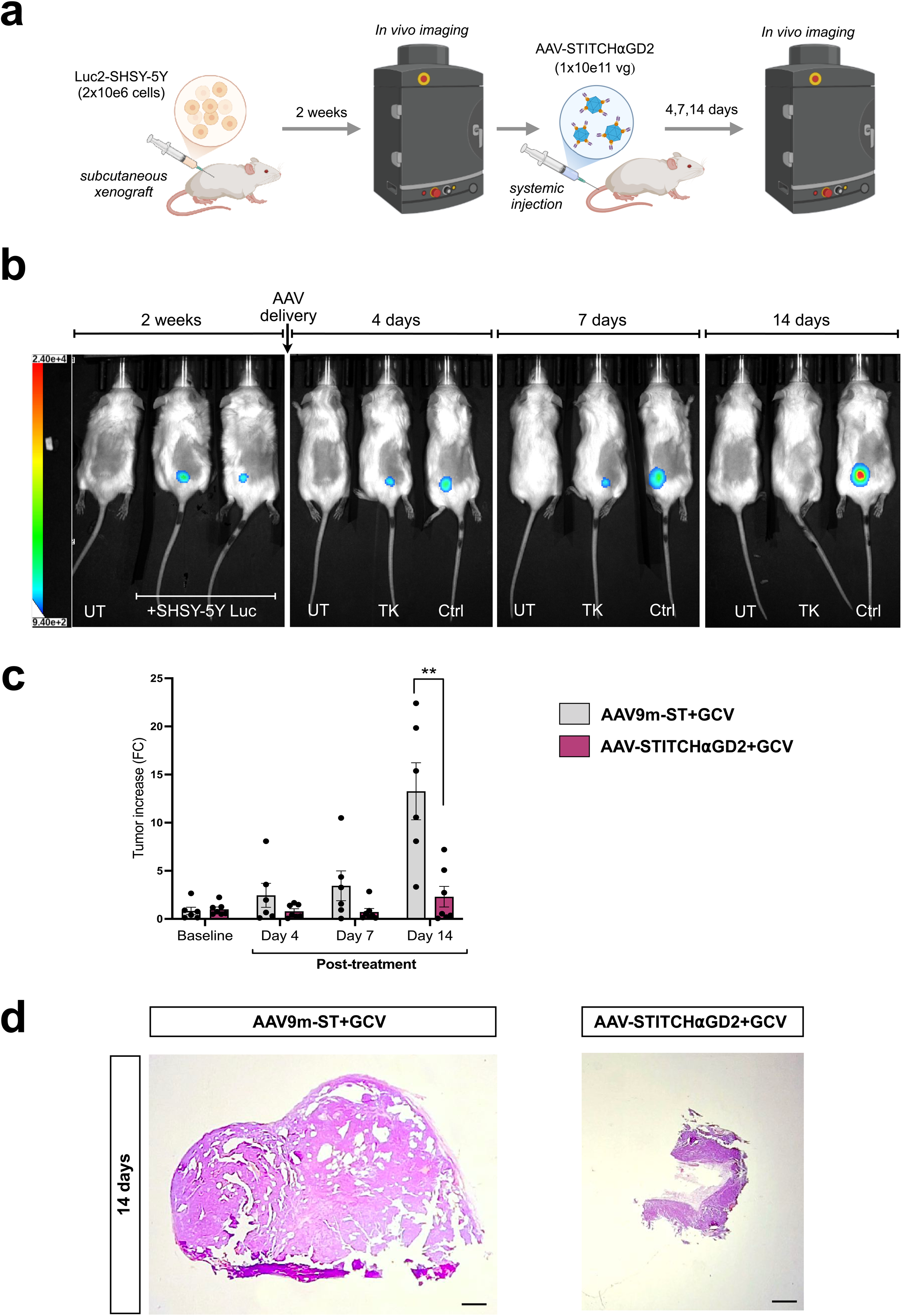
TK-expressing AAV-STITCHαGD2 inhibits the growth of subcutaneous tumor in xenograft mouse model. **a**. Scheme of the experimental protocol. In brief, 2x10e6 SH-SY5Y Luc+ cells were subcutaneously injected into adult NGS mice (xenograft model). Two weeks later, tumoral masses were determined by IVIS scan and AAV-STITCHαGD2 or AAV-ST (negative control) carrying HSV-TK transgene were inoculated in the tail vein using 1x10e11 vg/mouse. The following week, mice were daily injected with GCV and tumor growth was monitored by IVIS scan for 14 days. **b**. Representative bioluminescence images taken with IVIS Lumina of control and TK-treated xenograft mice acquired at different experimental time-points. **c**. Bar plot representing the quantification of the tumor growth evaluated though bioluminescence signal at 4 time points (baseline, 4-7-14 days post-treatment) in mice treated with AAV-STITCHαGD2 (n=7, purple) or AAV-ST (n= 6, gray). **d**. Representative haematoxylin and eosin (H&E) staining of subcutaneous tumors from control mice (AAV-ST) and mice treated with AAV-STITCHαGD2. Scale bar: 500 µm. Error bars represent SD. **p < 0.01. Two-way ANOVA with Sidak’s multiple comparisons correction.

### AAV-STITCHαGD2 dampens the detrimental effects of metastatic NB and inhibits patient-derived xenograft growth

Next, we ascertained the effects of the TK-expressing AAV-STITCHaGD2 in the pseudometastatic xenograft NB mouse model. Two weeks after intravascular administration of SH-SY5Y NB tumor cells, animals underwent MRI to identify those with comparable numbers of detectable hepatic metastases, which were then randomized into two groups to receive either AAV-STITCHαGD2 or the control (AAV9m-ST) viral treatment (2.5x10e10 vg) with GCV (Fig. 5a). Two weeks after viral infusion, a second MRI scan revealed a marked reduction in both the number and size of NB hepatic metastases in animals treated with AAV-STITCHαGD2 compared to controls (53,78 ± 6,34% of mean tumor growth inhibition) (Fig. 5b,c). The marked inhibition of metastatic growth was further confirmed by immunohistochemical visualization of Ki67+ proliferating cells in AAV-STITCHαGD2 treated liver tissues (Fig. 5d). Additionally, AAV-STITCHαGD2-treated mice showed a significant extension in survival compared to animals treated with the control vector (Fig. 5e).

**Figure 5.**
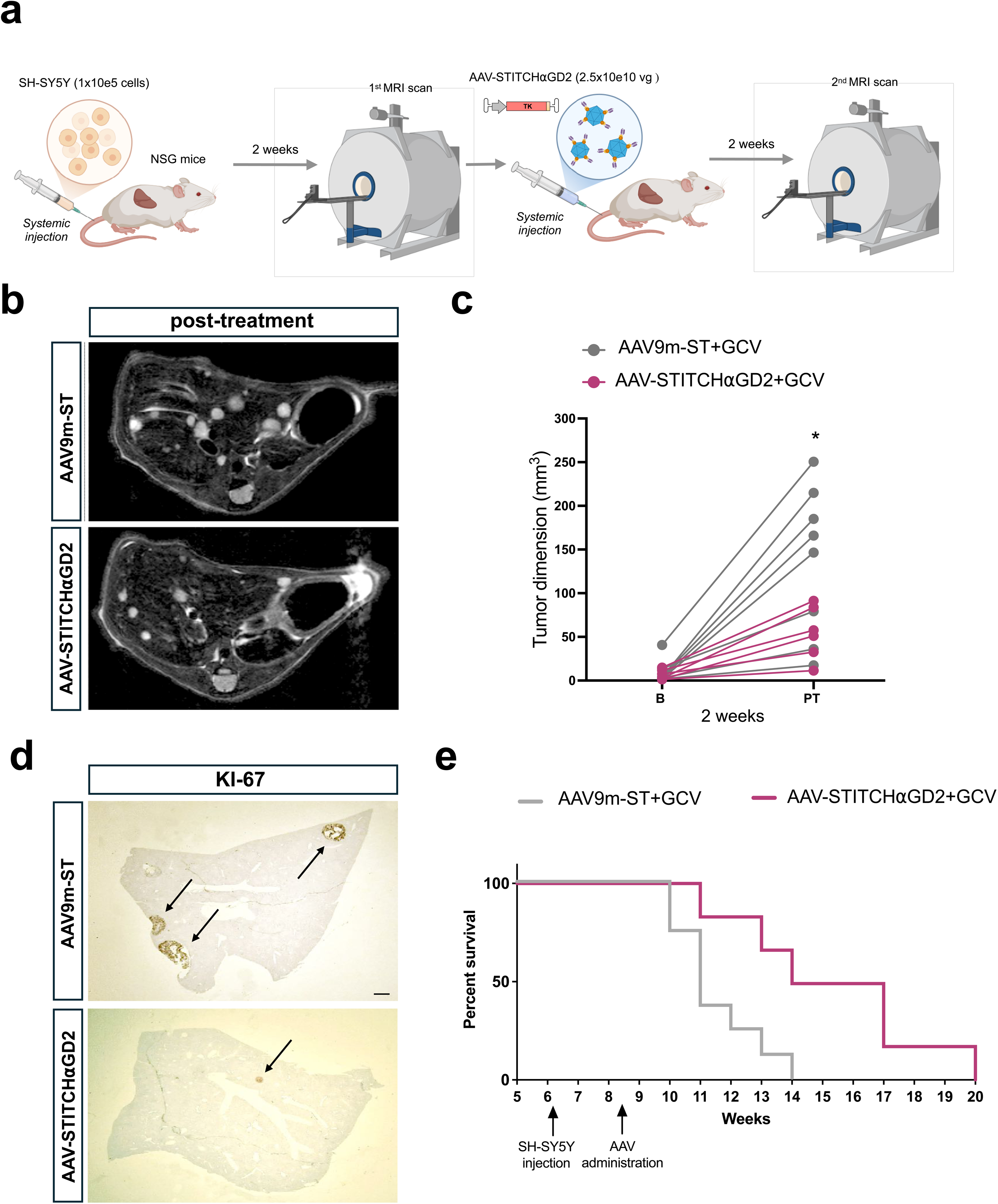
TK-expressing AAV-STITCHαGD2 inhibits neuroblastoma hepatic metastasis growth and significantly extends animal survival. **a**. Scheme of the experimental protocol. 1x10e5 SH-SY5Y cells were systemically injected into adult NGS mice (pseudometastatic model). Two weeks later, metastasis volume was determined by MRI and AAV-STITCHαGD2 or AAV-ST (negative control) carrying HSV-TK transgene were inoculated in the tail vein using 2.5x10e10 vg/mouse. The following week, mice were daily injected with GCV and after 10 days a second MRI scan was performed. **b**. Representative MRI images of liver metastasis before and after the treatment. **c**. Plot representing the volume of liver metastasis at the first (baseline) and at the second scan (post-treatment) in mice treated with AAV-STITCHαGD2 (n=6, purple) or AAV-ST (n= 8, gray). **d**. Representative staining for Ki67 staining in liver highlighting the size and localization of proliferating NB metastases (arrows) from control mice (AAV9m-ST) and mice treated with AAV-STITCHαGD2. Scale bar: 500 µm. **e**. Survival plot of mice treated with AAV-STITCHαGD2 (purple) or AAV-ST (gray) expressing HSV-TK + GCV. *p < 0.01. Two-way ANOVA with Sidak’s multiple comparisons correction and Mantel-Cox test for survival curves.

Finally, we tested the anti-tumoral activity of AAV-STITCHαGD2 in a patient-derived xenograft (PDX) mouse model. MYCN-amplified tumor explants from two patients with high-risk neuroblastoma (HR-NB; COG-N-424x and COG-N-480x) were obtained from the Childhood Cancer Repository and subcutaneously grafted into NSG mice (Fig. 6a). Two weeks later, animals received intravascular injections of AAV-STITCHαGD2 vectors expressing either Luc2 and TK, or Luc2 alone as a control (1x10e11 vg for each vector), allowing longitudinal intravital imaging of tumor growth following treatment (Fig. 6a). Longitudinal imaging at days 4 and 7 after treatment revealed a marked suppression of tumor growth in the AAV-STITCHαGD2-TK group compared to controls (Fig. 6b,c). In control mice, Luc2 signal progressively increased from day 0 to day 4, then plateaued by day 7, likely reflecting continued dilution of viral genomes in the proliferating tumor cells over time. In contrast, the Luc2 signal in the TK-treated group did not increase, indicating that luciferase-expressing tumor cells were efficiently eliminated following activation of the suicide gene (Fig. 6b,c). These findings confirm the potent anti-tumoral effects of AAV-STITCHαGD2 in suppressing malignant growth of primary human neuroblastoma tissue, highlighting its strong translational potential.

**Figure 6.**
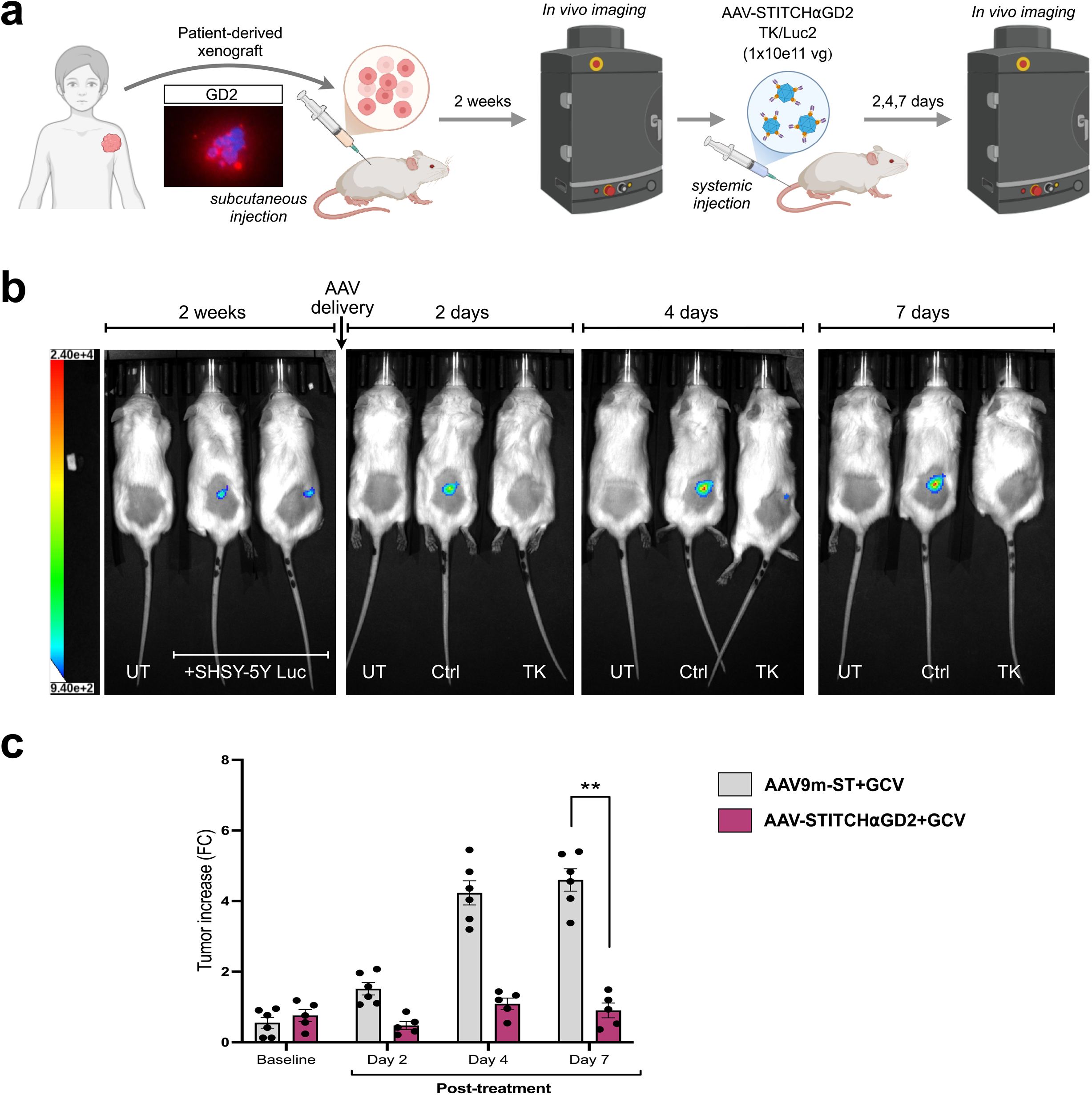
TK-expressing AAV-STITCHαGD2 inhibits the growth of subcutaneous tumor in PDX mouse model. **a.** Scheme of the experimental protocol. Patient-derived cancer cells positive for GD2 (Scale bar: 25 µm) were subcutaneously injected into adult NGS mice (PDX model). Two weeks later, AAV-STITCHaGD2 or AAV-ST (negative control), both carrying HSV-TK and luciferase transgenes, were administered via tail vein injection at a dose of 1x10e11 vg/mouse for each viral vector. Five days later, baseline tumor burden was assessed by IVIS imaging. Starting the following week, mice received daily injections of GCV, and tumor growth was monitored by IVIS imaging over 14 days. **b.** Representative bioluminescence images taken with IVIS Lumina of control and TK-treated PDX mice acquired at different experimental time-points. **c**. Bar plot representing the quantification of the tumor growth evaluated though bioluminescence signal at 4 time points (baseline, 2-7 days post-treatment) in mice treated with AAV-STITCHaGD2 (n=5, purple) or AAV-ST (n= 6, gray). Error bars represent SD. **p < 0.01. Two-way ANOVA with Sidak’s multiple comparisons correction.

## Discussion

We have developed a novel conjugation system that leverages the high affinity of the SpyTag/SpyCatcher interaction to enable efficient and specific covalent coupling of polypeptides to the surface of the AAV capsid. By incorporating the SpyTag into a permissive region of the infectivity-deficient AAV9-W503A vector, we created a capsid whose tropism can be reprogrammed by displaying different targeting molecules on its surface. Thus, this system serves as universal platform that can be exploited for any application by simply fusing the tropism-guiding molecule with the SpyCatcher. Importantly, we showed that the W503A mutation fully ablates the infection capability of the AAV9, generating a capsid without any residual off-target transduction. Tailored capsid modifications have driven the successful development of engineered AAV capsids with desirable new properties. Nevertheless, the broad, multi-tissue tropism of native AAV capsids remains a significant barrier to the development of safer clinical applications. Previous strategies to modify the capsid surface have included the conjugation of molecules via peptide fusions with the viral VP2 or VP3 proteins. However, only short peptide sequences can be incorporated without compromising capsid assembly or viral yield (19,20). More recently, site-specific incorporation of unnatural amino acids into the AAV capsid has enabled the covalent attachment of molecules through click chemistry (21,22). While promising, this method is limited by reduced AAV production titers and a labor-intensive, multi-step workflow. The AAV-STITCH system greatly broadens the potential for capsid coupling by enabling the attachment of any element that can be fused to the SpyCatcher, while maintaining high viral titers, offering exceptional versatility, and simplifying the overall viral production procedures.

Surprisingly, we found that the high coupling of the SpyCatcher-αGD2 onto the capsid is reducing the viral infectivity of GD2+ NB cell lines. We showed that this did not depend by the binding of the virus to the cell surface, but by the subsequent steps of viral transduction. This impairment aligns with previous observations showing that an excessive coupling of FITC molecules onto the AAV2 capsid reduced its infectivity in HEK293 cells (23). It is likely that an excessive molecule display masks the recognition of AAV co-receptors on the membrane surface or along its intracellular route. More studies are warranted to define which exact step in the AAV transduction cycle is impaired by the excessive loading of scFv moieties on the capsid surface.

We prioritize the coupling of αGD2 moieties to develop a valuable anti-tumor agent targeting NB. GD2 is a highly and consistently expressed antigen in this tumor type, a feature that has already been successfully exploited in antibody-based and CAR T cell therapies. However, these immunotherapies face limitations due to the immunosuppressive tumor microenvironment and potential off-target effects. Importantly, the tropism and therapeutic efficacy of our viral approach are not affected by the tumor’s immune status. Although GD2 is also expressed in peripheral neurons, the AAV-STITCHαGD2-TK system poses no risk to these cells, as the TK/GCV suicide gene system is selectively active in proliferating cells. Furthermore, the vector design offers the flexibility to incorporate additional regulatory elements, such as cell cycle–specific promoters or cell type–specific microRNA target sequences, to further enhance the precision of TK expression. The strong anti-tumoral effects of TK/GCV treatment relies also by its significant bystander effect since the phosphorylated GCV is passively transferred to surrounding TK-non-expressing tumor cells inducing their cell death (24). Moreover, TK/GCV-mediated cell death might trigger inflammation and T cell recruitment harnessing the immune response against the tumor (25). Therefore, AAV-STITCHαGD2-TK not only provides a complementary strategy to existing immunotherapies but can enhance their anti-tumoral effects by modulating the tumor microenvironment. A current limitation of AAV-based therapies is that they can typically be administered only once, and only to seronegative individuals. However, since neuroblastoma primarily affects children, a population with a lower prevalence of pre-existing AAV immunity, it is estimated that approximately 80% of patients would be eligible (26). Treated patients will develop AAV neutralizing antibodies that will exclude a second viral treatment, representing a current limitation for this treatment. However, new advances in the generation of AAV capsids with reduced immunogenicity and clinical strategies to overcome pre-existing antibodies will provide new opportunities for repetitive viral administration in the near future (27-30).

In summary, we have developed AAV-STITCH, a rapid and versatile conjugation system that enables the targeted coupling of polypeptides to the surface of an infectivity-deficient AAV9 capsid, facilitating cell-type-specific viral transduction. This platform addresses longstanding challenges in the field by improving AAV selectivity and gene transduction efficiency, thereby expanding the potential of AAV-based therapies in clinical applications. As a compelling proof-of-concept, we engineered the AAV-STITCHαGD2 system, which functions as a potent anti-tumor agent capable of inhibiting the growth of NB xenografts derived from both established tumor cell lines and primary tumor tissues. This study introduces and validates a radically new viral anti-tumor strategy with unique, complementary advantages over existing therapeutic options. Consequently, this approach is well positioned as an adjuvant treatment to broaden and strengthen the therapeutic arsenal available for patients with neuroblastoma.

## Acknowledgements

We thank M. Brenner (Baylor College of Medicine, Houston, TX) for the scFv-anti-GD2 sequence and all members of the Broccoli’s lab for helpful discussion. We are grateful to the Childhood Cancer Repository and Alex’s Lemonade Stand Foundation for providing patient-derived NB tissues. We acknowledge the FRACTAL and ALEMBIC core facilities for expert supervision in flow-cytometry and confocal imaging, respectively. Image items used in this work were created in BioRender (https://BioRender.com). This work was supported by the PNRR-National Center for Gene Therapy and Drugs Based on RNA Technology (project #CN00000041; B83C22002860006) and EU NRRP “D34Health” (project #PNC0000001; CUP B53C22006100001) to V.B.

## Author contributions

V.B. and M.L. conceptualized the project. M.L., S.G.G., S.P., F.M., T.C., A.D.P., S.M., A.I., E.V. and T.R. conducted the experiments. A.S. contributed to the in vivo imaging analyses. F.P. contributed to the design of the in vivo experiments. V.B. and M.L. wrote the paper with input from all authors.

## Competing financial interests

L.M., S.P. and V.B. are inventors of filed patents based on the work published here. The other authors declare no competing interests.

## Methods

### Plasmids

The SpyTag peptide sequence was inserted into the AAV9-A503W plasmid between amino acids 453 and 454 using overlap extension PCR (OEP), exploiting the NheI and AgeI restriction sites. The SpyCatcher peptide sequence, fused to the scFv-anti-GD2 (a kind gift from M. Brenner), was synthesized by Azenta Genewiz and subcloned into pcDNA3.1 plasmid (Addgene) with a V5 tag inserted between the two peptides and a His-tag added at the C-terminus of the construct. scAAV-CMV vectors carrying the ZsGreen, Luciferase, and HSV-TK transgenes were generated by amplifying the respective coding sequences (CDS) and cloning them into the scAAV-GFP backbone (Addgene #32396) using BamHI and NotI restriction sites. Finally, a bicistronic lentiviral vector (Luciferase-IRES-Puro) was generated by amplifying the Luciferase CDS and cloning it into the Lv-Ef1a-IRES-Puro vector (Addgene #85132) using BamHI and EcoRI restriction sites.

### Post-production coupling protocol

#### AAV production and purification

AAV replication-incompetent, recombinant viral particles were produced in 293T cells, cultured in Dulbecco Modified Eagle Medium high glucose (Sigma-Aldrich) containing 10% fetal bovine serum (Sigma-Aldrich), 1% non-essential amino acids (Gibco), 1% sodium pyruvate (Sigma-Aldrich), 1% glutamine (Sigma-Aldrich) and 1% penicillin/streptomycin (Sigma-Aldrich). Cells were split every 3-4 days using Trypsin 0.25% (Sigma-Aldrich). Recombinant viral particles were produced in 293T cells by polyethylenimine (PEI, Polyscience) co-transfection of three different plasmids: transgene-containing plasmid (scAAV), packaging plasmid for rep and cap (AAV9-W503A and AAV9-W503A-SpyTag at different ratios) genes, pHelper (Agilent). The cells and supernatant were harvested at 120 hrs. Cells were lysed in hypertonic buffer (40mM Tris, 500mM NaCl, 2mM MgCl2, pH=8) containing 100U/ml Salt Active Nuclease (SAN, Arcticzymes) for 1h at 37°C, whereas the viral particles present in the supernatant were concentrated by precipitation with 8% PEG8000 (Polyethylene glycol 8000, Sigma-Aldrich) and then added to supernatant for an additional incubation of 30min at 37°C. In order to clarify the lysate cellular debris were separated by centrifugation (4000g, 30min). The viral phase was isolated by iodixanol step gradient (15%, 25%, 40%, 60% Optiprep, Sigma-Aldrich) in the 40% fraction and concentrated in PBS (Phosphate Buffer Saline) with 100K cut-off concentrator (Amicon Ultra15, MERCK-Millipore). Virus titers were determined using AAVpro© Titration Kit Ver2 (TaKaRa).

#### AntiGD2-SC protein purification

293T cells were cultured in DMEM (Sigma) supplemented with 10 % FBS (Gibco) and 4 mM L-Gln (Sigma). To express αGD2-SpyCatcher (αGD2-SC) fusion protein, HEK293 cells were transiently transfected with the plasmid encoding histidine-tagged αGD2-SC, using PEI (Polyscience). After 24 h transfected cells were washed with phosphate buffered saline (PBS) and cultured in the serum-free Pro293a-CDM media (Lonza) supplemented with 4mM L-Gln for an additional 72 h. Conditioned media was harvested, centrifuged to remove cell debris, and concentrated ten times using centrifugal filter units with a 10kDa cutoff (Amicon, Merk Millipore). Concentrated conditioned media was dialyzed in 20 mM sodium phosphate, 0.5 M NaCl, pH 7.4. His-tagged αGD2-SC was purified from the conditioned media by affinity chromatography on Ni Sepharose 6 Fast Flow (Sigma). Ni Sepharose-adsorbed αGD2-SC was eluted in 20 mM sodium phosphate, 0.5 M NaCl, pH 7.4 containing 500 mM imidazole and dialyzed in PBS. Purified αGD2-SC was analyzed by 10% SDS-PAGE and by immunoblot. In immunoblot experiments αGD2-SC was detected using an anti-V5 antibody (#R960-25, Thermo Scientific).

### Co-expression coupling protocol

AAV replication-incompetent, recombinant viral particles were produced as described before. For targeted capsids, 1.5 μg of the SpyCatcher-αGD2 plasmid in 10 cm dish was added together with the other three plasmids. Mosaic capsids were generated by co-transfecting rep/cap plasmids (AAV9-W503A) containing or lacking the SpyTag insertion at defined ratios (1:3 and 1:7).

### Western Blot

Western Blots analysis was performed on purified AAV vector preparations using a dose of 1x10^9^ vg. Proteins were denaturated at 95°C in Laemmli buffer containing SDS, β-mercaptoethanol and dithiothreitol (DTT). Proteins were then separated via SDS-PAGE using 8% polyacrylamide gel and then transferred to nitrocellulose membranes. Membranes were incubated overnight at 4°C with the following primary antibodies in 1X PBST with 5% w/v nonfat dry milk: mouse anti-AAV VPs (1:250; Arp1), mouse anti-V5 (1:1000; ThermoFisher Scientific). Subsequently, membranes were incubated with the corresponding horseradish peroxidase (HRP)-conjugated secondary antibodies (1:10000; Dako). The signal was then revealed with a chemiluminescence solution (ECL reagent, RPN2232; GE Healthcare) and detected with the ChemiDoc imaging system (BioRad).

### Cell cultures

Human neuroblastoma cell lines were cultured in plastic-adherence condition in RPMI 1640 medium (Roswell Park Memorial Institute 1640 Medium, Gibco) containing 10% fetal bovine serum (FBS, Sigma-Aldrich), 1% Pen/Strept (Sigma-Aldrich), 2mM Glutamine (Sigma-Aldrich), 1% non-essential amino acids (MEM NEAA, ThermoFisher Scientific), 1% sodium pyruvate solution (Sigma-Aldrich) and passaged twice a week using Trypsin-EDTA solution (Sigma-Aldrich). All cell cultures were kept in humidified atmosphere of 5% CO2 at 37°C under atmospheric oxygen conditions.

### *In vitro* AAV transduction and immunofluorescence

25.000 cells per well were plated on glass coverslips in 500 μL of media in a 24-well plate. The following day, cells were transduced with AAV-ST or AAV-STITCH variants at a Multiplicity of Infection (MOI) of 40.000 in case of low titer testing and with an MOI of 400.000 in case of high titer testing. 24 hours after transduction the media was changed for fresh media.

72 hours post transduction cells were fixed with ice-cold 4% paraformaldehyde (PFA) for 30 min at 4°C, washed with PBS (3×) and then incubated O/N with primary antibody: mouse anti-GD2 (1:300, BD Biosciences). The following day, upon wash with PBS (3×), cells were incubated for 1 h at RT in blocking solution with DAPI and with Alexa Fluor-594 anti-mouse secondary antibodies. After PBS washes (3×), cells were mounted with fluorescent mounting medium (Dako). Images were captured with a Nikon Eclipse 600 fluorescent microscope.

### Immunohistochemistry

Subcutaneous tumor and metastatic liver tissues were harvested, fixed in 4% PFA for 24–48 hours at 4°C, and subsequently processed for paraffin embedding. Formalin-fixed paraffin-embedded (FFPE) tissues were sectioned at 5 μm using a microtome and mounted onto glass slides. For histological evaluation, sections were deparaffinized in xylene, rehydrated through a graded ethanol series, and stained with haematoxylin and eosin (H&E) following standard protocols. For immunohistochemical analysis, antigen retrieval was performed by heating the slides in citrate buffer (pH 6.0), followed by blocking in appropriate serum. Sections were incubated overnight at 4°C with an anti-Ki67 primary antibody (Abcam), then with a biotinylated secondary antibody and an avidin-biotin complex (ABC, Abcam) detection system. Signal was visualized using diaminobenzidine (DAB, Sigma). After staining, slides were dehydrated, cleared in xylene, and cover slipped using a permanent mounting medium. Images were acquired using a brightfield microscope.

### Organotypic Brain Cultures

Organotypic brain slice cultures were prepared from adult C57BL/6 wild-type mice. Brains were extracted and sectioned into 300 μm-thick coronal slices using a vibratome. Slices were placed onto porous membrane inserts within 6-well culture plates and maintained in culture medium composed of Neurobasal (Thermo Fisher), 1× B27 supplement (Thermo Fisher), 1× 30% glucose (Sigma), and 1× penicillin/streptomycin and glutamine (Sigma). Slices were then transduced overnight with AAV-STITCHαGD2 at a dose of 2 × 10¹⁰ vg/well. The following day, slices were fixed in 4% paraformaldehyde (PFA), embedded in OCT compound, sectioned at 30 μm, and stained with an anti-ZsGreen antibody. Images were captured with Maving RS-G4 Confocal and Nikon Eclipse 600 fluorescent microscope.

### Virus Binding Assays

Virus binding assays were performed as described previously (31). In brief, AAVs were added to SH-SY5Y at 40.000 MOI for 1 hr at 4°C. After, 3x PBS washes, cells were lysed in 1% NP-40 buffer, and viral DNA was extracted and quantified using AAVpro© Titration Kit Ver2 (TaKaRa).

### Total DNA isolation for vector copy number

Total DNA was isolated from primary neurons and animal tissues (cortex and liver) using the QIAamp DNA Micro (QIAGEN). The quantification of vector transgene expression was calculated by qRT-PCR using primers on ZsGreen sequence described in 14. The DNA levels were normalized against an amplicon from a single-copy human gene, *LMNB2*, amplified from genomic DNA.

### *In vitro* cytotoxicity assay

Citotoxicity was assessed via LIVE/DEAD™ Viability/Cytotoxicity Kit (ThermoFisher). Briefly, 25.000 SH-SY5Y cells were plated on glass coverslips in 500 μL of media in a 24-well plate. The following day, cells were transduced with the viral vector (AAV-STITCHaGD2) followed by a daily exposition to the prodrug 8 ug/ml Ganciclovir (Sigma-Aldrich) for 72 h or to DMSO as negative control. Then staining solution was prepared by adding 5 µL of 4 mM Calcein AM and 20 µL of 2 mM Ethidium Homodimer III (EthD-III) to 10 mL of PBS. Cell media was removed before addition of 200ul/well of staining solution, incubated for 30 minutes at RT. Then, cells were mounted with fluorescent mounting medium (Dako). Images were captured with a Nikon Eclipse 600 fluorescent microscope.

### *In vivo* neuroblastoma models generation

#### Pseudometastatic model

SH-SY5Y cells were resuspended in sterile PBS at a concentration of 1x10e6 cells/ml and then 100 µl (100.000 cells) were injected into the tail vein of NSG mice (NOD.Cg-*Prkdc^scid^ Il2rg^tm1Wjl^*/SzJ) using 0,3 ml syringes. The extent of liver metastasis was then asses through MRI imaging at 2- and 3-weeks post injection for metastatic burden evaluation.

#### Subcutaneous model

3x10e5 SH-SY5Y cells were seeded in a well of a 6 multi-well plate and transduced with LV-Luc-ires-Puro (10e8 IFU/ml). After 72h of puromycin selection, we expanded the cells in a T25 flask. SH-SY5Y-Luc+ cells were then resuspended in a 50% Matrigel (Matrigel growth factor reduced, Corning) solution in sterile PBS at a concentration of 2x10e7 cells/ml. Then 100 µl per mouse of solution were subcutaneously injected in the right flank of NSG mice (NOD.Cg-*Prkdc^scid^ Il2rg^tm1Wjl^*/SzJ) using 0,3 ml syringes. Evaluation of tumor burden progression occurred through BLI imaging (IVIS spectrum) at multiple timepoints from the second week post injection onward trough intraperitoneal luciferin injection.

#### Patient-Derived Xenograft (PDX) Model

Patient-derived cells were kindly donated by Childhood Cancer Repository (https://www.cccells.org/). 5x10e6 cells were injected subcutaneously in NSG mice (NOD.Cg-*Prkdc^scid^ Il2rg^tm1Wjl^*/SzJ) using 0,3 ml syringes. After two weeks, mice were injected intravenously with AAV-STITCHaGD2 expressing Luciferase to evaluate tumor burden progression through BLI imaging (IVIS spectrum) at multiple timepoints onward trough intraperitoneal luciferin injection.

### *In vivo* AAV-STITCHaGD2 transduction

Pseudometastatic neuroblastoma mouse model was generated as described in previous section. After two weeks from SH-SY5Y cell injection, mice were injected systemically (tail vein) with ZsGreen encoding AAV-STITCHaGD2 vector (1x10e11 vg/mouse). 72 h after tail vein injection of the vector, mice were sacrificed and transcardially perfused in saline solution. The following organs were harvested and post-fixed in 4% PFA in PBS for 48h: liver (containing both stroma and metastases), heart, kidney, muscle, cortex and cerebellum. After fixation, organs were soaked in cryoprotective solution (30% sucrose in PBS) to prepare samples for cryostat sectioning and anti-ZsGreen immunofluorescence analysis.

### Tissue immunofluorescence staining

Tissues were sectioned using cryostat after optimal cutting temperature compound (OCT) embedding in dry ice. 50μm-thick histological slides of each collected organ were mounted on a glass slide and rinsed in PBS and were incubated with 10% donkey serum (Sigma-Aldrich) and 3% Triton X-100 (Sigma-Aldrich) for 1 hr at RT to saturate the unspecific binding site before the overnight incubation at 4°C with the primary antibody (diluted in the blocking solution).

Upon wash with PBS (3×), sections were incubated for 1 h at RT in blocking solution with DAPI (1:1000, Sigma-Aldrich) and with Alexa Fluor-488 anti-rabbit secondary antibody (1:1000, ThermoFisher Scientific). After PBS washes (3x), sections were mounted with fluorescent mounting medium (DAKO). Images were captured with a Nikon Eclipse 600 fluorescent microscope. Tissues were stained with the following primary antibody: rabbit Living Colors® Full-Length ZsGreen Polyclonal Antibody (1:500; Takara Bio).

### *In vivo* cytotoxicity essays

Pseudometastatic and subcutaneous xenograft mouse models were injected systemically (tail vein) with AAV-STITCHaGD2 expressing HVS-TK two weeks after SH-SY5Y inoculation. Then all mice were intraperitoneally injected with 1 mg/kg of Ganciclovir (Sigma-Aldrich) for seven consecutive days. At the end of the treatment, tumor burden evaluation was performed either through MRI (pseudometastatic model) or IVIS BLI (subcutaneous model) to assess changes in tumor growth between experimental groups. MRI imaging analysis were performed using “MIPAV” software to quantify both metastasis number and dimension (mm^3^). BLI imaging analysis were performed through “Aura” software, for quantification of photon intensity from the tumor burden.

PDX mouse models were injected with AAV-STITCHaGD2 expressing HVS-TK one week after Luciferase injection. Then mice were intraperitoneally injected with 1 mg/kg of Ganciclovir for seven days and tumor burden evaluation was performed through IVIS BLI as described previously.

### Statistics

Values are expressed as mean ± standard deviation as indicated. All statistical analysis was carried out in GraphPad Prism 8.0, using *t*-Test, ANOVA-one way with Tukey’s post hoc test, Two-way ANOVA with Sidak’s multiple comparisons correction and Mantel-Cox test for survival curves. Number of replicates for statistical testing were indicated in corresponding figure legends.

## Supplementary Figures

**Supplementary Figure 1.**
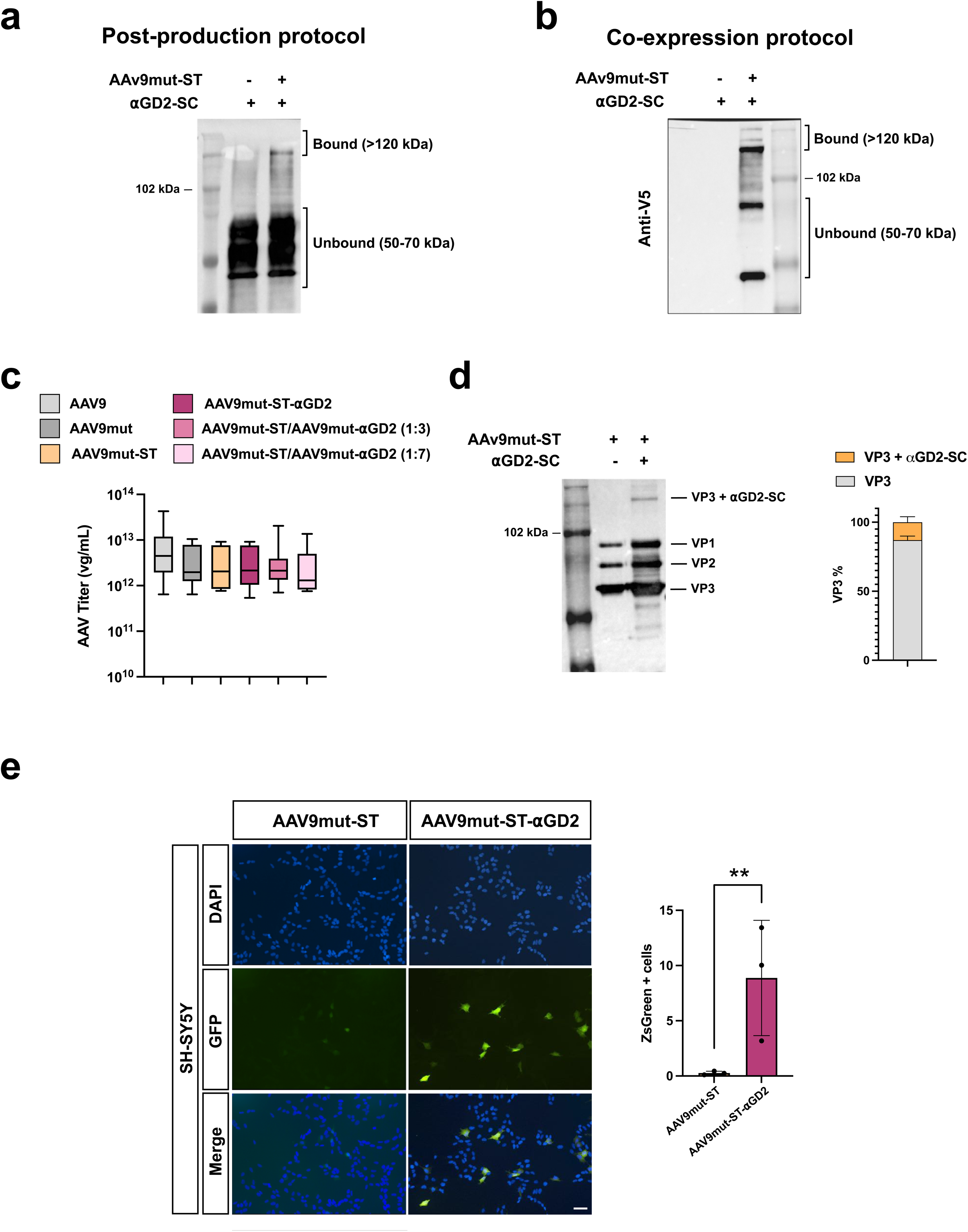
Optimization of AAV-STITCH production and characterization of capsid conjugation efficiency a–b. Representative anti-V5 Western blot showing the detection of SpyCatcher fused to the anti-GD2 scFv (⍺GD2-SC) alone or co-expressed with AAV9-mut-ST either post-production (**a**) or during viral production (**b**). **c**. Box plot showing the viral genome titers, assessed by RT-qPCR, from multiple independent productions of AAV9, AAV9mut, AAV9mut-ST, and 1:3 and 1:7 mosaic preparations (n > 3 experimental replicates per group). **d**. Left: Anti-AAV Western blot showing the capsid composition of AAV9mut-ST alone or conjugated to ⍺GD2-SC. Right: Densitometric quantification of the proportion of unconjugated VP3 (gray) and VP3 conjugated with ⍺GD2-SC (orange). **e**. On the left, immunofluorescence using anti-ZsGreen antibody on SH-SY5Y cells transduced with control AAV9mut-ST alone or couple with ⍺GD2. Scale bar: 100 µm. On the right, bar graph showing the quantification of ZsG+ cells over total cells in the different experimental conditions. n=3 independent biological replicates. Error bars represent SD. **p < 0.01; t-test.

**Supplementary Figure 2.**
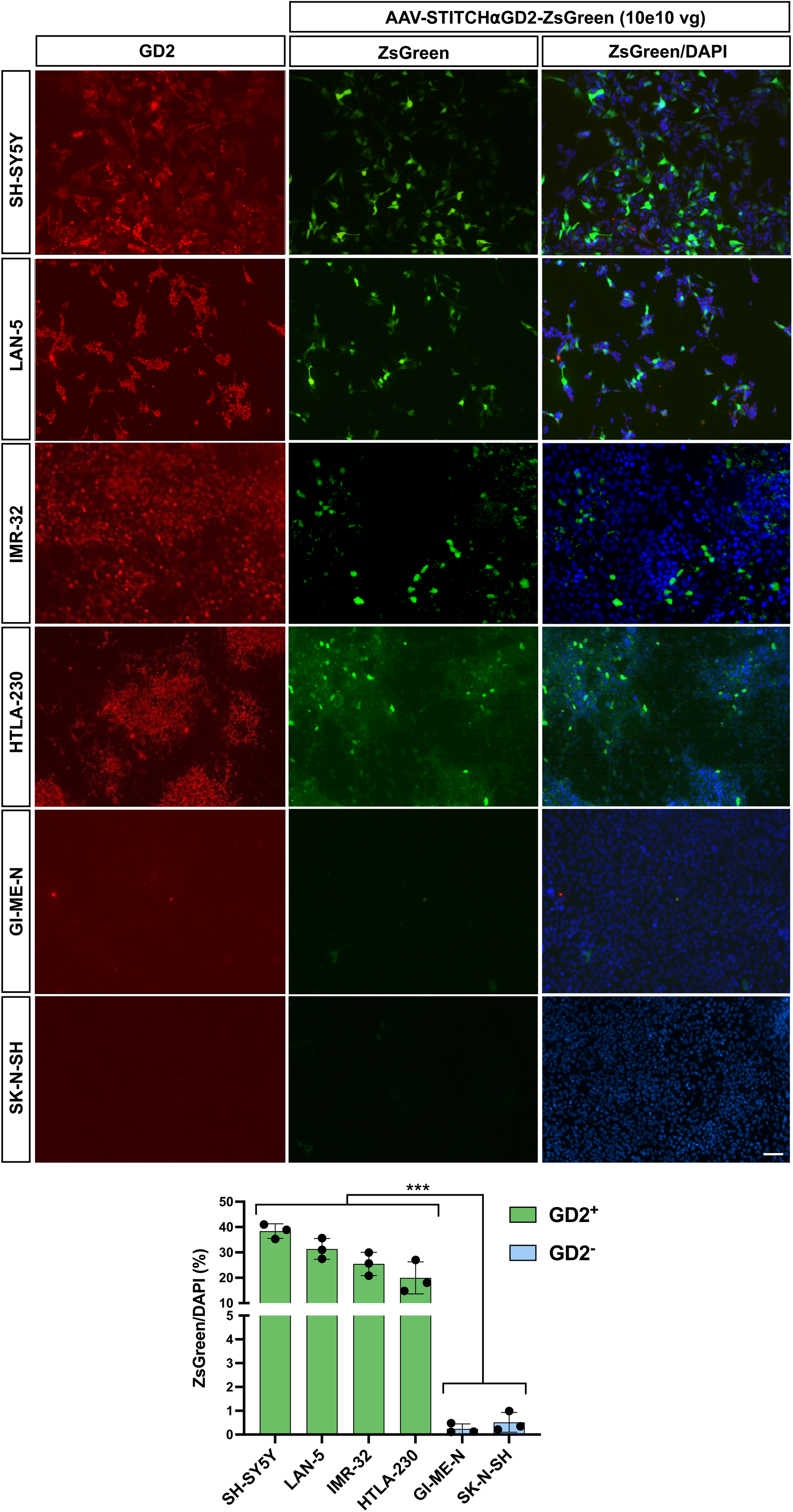
Specificity and efficiency of AAV-STITCHaGD2 in neuroblastoma cell lines. Above, representative immunofluorescence for anti-GD2 and anti-ZsGreen in GD2+ and GD2-neuroblastoma cell lines. Scale Bar: 100 µm. Below, bar graph showing the quantification of ZsG+ cells over total cells in the different neuroblastoma cell lines transduced with AAV-STITCHαGD2 carrying the ZsGreen transgene. n = 3 biological replicates. Error bars represent SD. ***p < 0.001. One-way ANOVA with Tukey’s multiple comparisons correction.

**Supplementary Figure 3.**
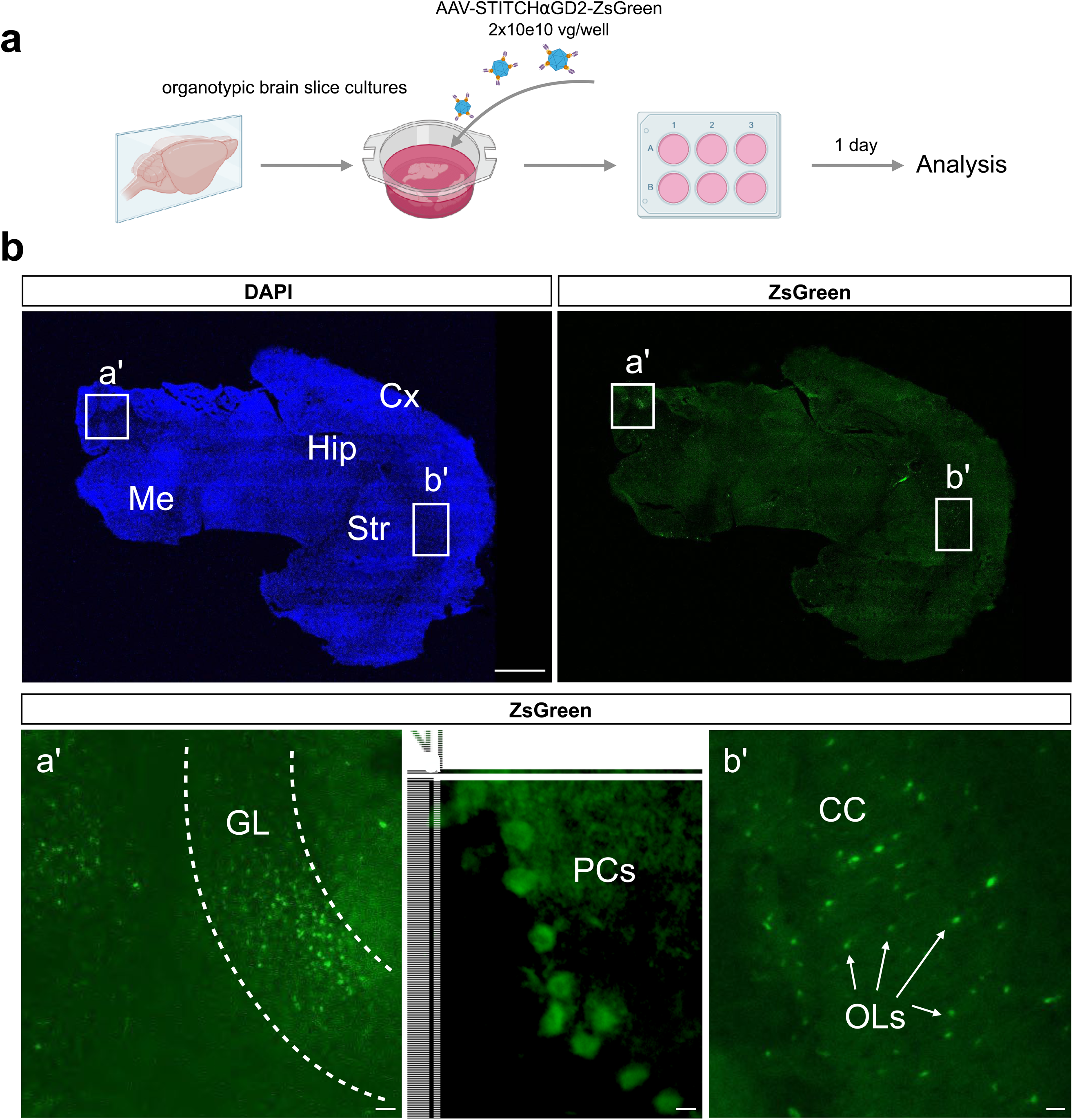
Transduction profile of AAV-STITCHaGD2 in organotypic murine brain slices. **a.** Schematic of the experimental protocol. Briefly, murine organotypic brain slices were infected with 2x10e10 vg/well of AAV-STITCHaGD2 carrying the ZsGreen transgene. The following day, slices were processed, and the transduction profile was analyzed by immunofluorescence. **b**. Representative immunofluorescence images of a sagittal brain section transduced with AAV-STITCH, showing DAPI staining on the left and anti-ZsGreen staining on the right. Scale bar: 1000 µm. **a′**. Left: Representative higher magnification image of the cerebellum, showing transduced cells in the granular layer (GL). Scale bar: 100 µm. Right: Higher magnification image of cerebellar Purkinje cells (PCs). Scale bar: 100 µm. **b′**. Representative higher magnification image showing transduced oligodendrocytes (OLs) in the corpus callosum (CC). Scale bar: 100 µm.

**Supplementary Figure 4.**
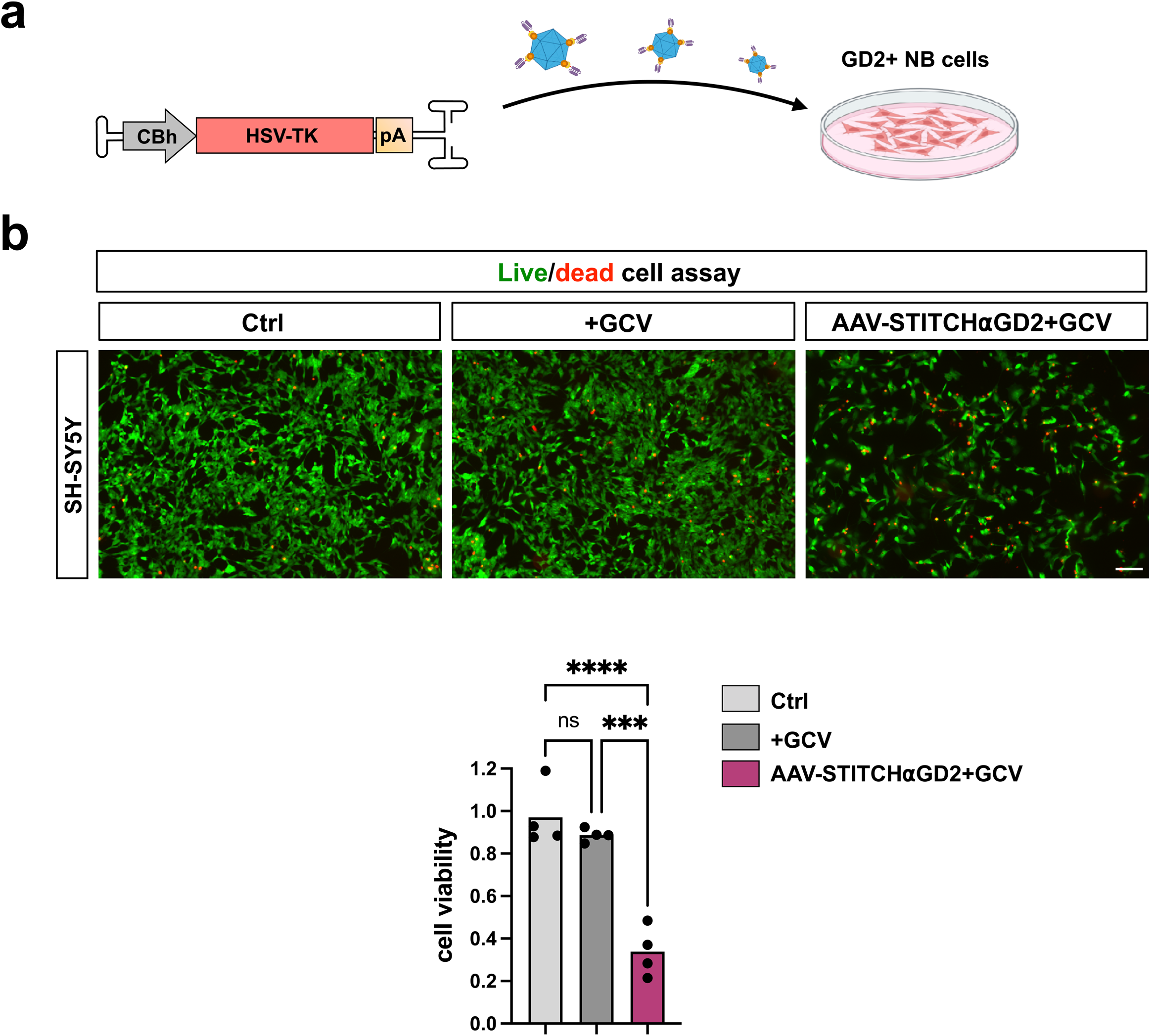
Evaluation of HSV-TK/GCV-mediated cytotoxicity in GD2+ neuroblastoma cells. **a.** Schematic of the experimental protocol used to assess HSV-TK activity in GD2+ SH-SY5Y neuroblastoma cells. **b**. Top: Immunofluorescence images using the Live/Dead Assay Kit to evaluate cell death in SH-SY5Y cells transduced with AAV-STITCH⍺GD2 and treated with Ganciclovir (GCV), compared to non-infected controls with or without GCV treatment. Scale bar: 100µm. Bottom: Bar graph showing quantification of cell viability across the three experimental conditions. n=3 independent biological replicates. Error bars represent SD. ***p<0.01. One-way ANOVA with Tukey’s multiple comparisons correction.

## References

1. Matthay KK, et al. Neuroblastoma. Nat Rev Dis Primers 2, 16078 (2016).

2. - Qiu B & Matthay KK. Advancing therapy for neuroblastoma. Nat.Rev.Clin.Oncol. 19, 515–533 (2022).

3. Yu AL et al. Anti GD2 antibody with GM-CSF, interleukin-2, and isotretinoin for neuroblastoma. N Engl J Med. 363, 1324–1334 (2010).

4. Mora J, et al. Desensitizing the autonomic nervous system to mitigate anti-GD2 monoclonal antibody side effects. Front Oncol. 14, 1380917 (2024).

5. Pule MA et al. Virus-specific T cells engineered to coexpress tumor-specific receptors: persistence activity in individuals with neuroblastoma. Nat Med.14, 1264–1270 (2008).

6. Straathof K et al. Antitumor activity without on-target off-tumor toxicity of GD2-chimeric antigen receptor T cells in patients with neuroblastoma. Sci Transl Med. 12, eabd6169 (2020).

7. Del Bufalo F, et al. GD2-CART01 for relapsed or refractory high-risk neuroblastoma. N Engl J Med. 388, 1284–1295 (2023).

8. Kinkhabwala A et al. MACSima imaging cyclic staining (MICS) technology reveals combinatorial target pairs for CAR T cell treatment of solid tumors. Sci Rep. 12, 1911 (2022).

9. Wang JH, Gessler DJ, Zhan W, Gallagher TL & Gao G. Adeno-associated virus as a delivery vector for gene therapy of human diseases. Signal Transduct Target Ther. 9, 78 (2024).

10. Deverman BE, Ravina BM, Bankiewicz KS, Paul SM & Sah DWY. Gene therapy for neurological disorders: progress and prospects. Nat Rev Drug Discov. 17, 641–659 (2018).

11. Haery L, et al. Adeno-Associated Virus Technologies and Methods for Targeted Neuronal Manipulation. Front Neuroanat. 13, 93 (2019).

12. Zakeri B, et al. Peptide tag forming a rapid covalent bond to a protein, through engineering a bacterial adhesin. Proc Natl Acad Sci U S A. 109, E690–7 (2012).

13. Chen X et al. Functional gene delivery to and across brain vasculature of systemic AAVs with endothelial-specific tropism in rodents and broad tropism in primates. Nat Commun. 14, 3345 (2023).

14. Giannelli SG, et al. New AAV9 engineered variants with enhanced neurotropism and reduced liver off-targeting in mice and marmosets. iScience 27, 109777 (2024).

15. McCarty DM. Self-complementary AAV vectors; advances and applications. Mol Ther. 16, 1648–1656 (2008).

16. Lammie G, Cheung N, Gerald W, Rosenblum M, Cordoncardo. Ganglioside gd(2) expression in the human nervous-system and in neuroblastomas - an immunohistochemical study. Int J Oncol. 3, 909–915 (1993).

17. Marconi S, et al. Expression of gangliosides on glial and neuronal cells in normal and pathological adult human brain. J Neuroimmunol. 170, 115–121 (2005).

18. Pastorino F, et al. Tumor regression and curability of preclinical neuroblastoma models by PEGylated SN38 (EZN-2208), a novel topoisomerase I inhibitor. Clin Cancer Res. 16, 4809–4821 (2010).

19. Münch RC, et al. Displaying high-affinity ligands on adeno-associated viral vectors enables tumor cell-specific and safe gene transfer. Mol Ther. 21, 109–118 (2013).

20. Münch RC, et al. Off-target-free gene delivery by affinity-purified receptor-targeted viral vectors. Nat Commun. 6, 6246 (2015).

21. Katrekar D, Moreno AM, Chen G, Worlikar A & Mali P. Oligonucleotide conjugated multi-functional adeno-associated viruses. Sci Rep. 8, 3589 (2018).

22. Puzzo F et al. Aptamer-programmable adeno-associated viral vectors as a novel platform for cell-specific gene transfer. Mol Ther Nucleic Acids 31, 383–397 (2023).

23. Mével, M. et al. Chemical modification of the adeno-associated virus capsid to improve gene delivery. Chem. Sci. 11, 1122–1131 (2019).

24. Oishi T et al. Efficacy of HSV-TK/GCV system suicide gene therapy using SHED expressing modified HSV-TK against lung cancer brain metastases. Mol Ther Methods Clin Dev. 26, 253–265 (2022).

25. Umegaki S, Shirota H, Kasahara Y, Iwasaki T & Ishioka C. Distinct role of CD8 cells and CD4 cells in antitumor immunity triggered by cell apoptosis using a Herpes simplex virus thymidine kinase/ganciclovir system. Cancer Sci. 114, 3076–3086 (2023).

26. Calcedo R. et al. Adeno-associated virus antibody profiles in newborns, children, and adolescents. Vaccine Immunol. 18, 1586–1588 (2011).

27. Chan YK et al. Engineering adeno-associated viral vectors to evade innate immune and inflammatory responses. Sci Transl Med. 13, eabd3438 (2021).

28. Meliani A, et al. Antigen-selective modulation of AAV immunogenicity with tolerogenic rapamycin nanoparticles enables successful vector re-administration. Nat Commun. 9, 4098 (2018).

29. Leborgne C et al. IgG-cleaving endopeptidase enables in vivo gene therapy in the presence of anti-AAV neutralizing antibodies. Nat Med. 26, 1096–1101 (2020).

30. Ronzitti G, Gross DA & Mingozzi F. Human Immune Responses to Adeno-Associated Virus (AAV) Vectors. Front Immunol. 11, 670 (2020).

31. Nonnenmacher M & Weber T. Adeno-associated virus 2 infection requires endocytosis through the CLIC/GEEC pathway. Cell Host Microbe. 10, 563–576 (2011).

