## Supplementary figures and images for "Targeted and efficient AAV therapy for neuroblastoma via direct capsid-antibody coupling"

### Supplementary Figures

#### Supplementary Figure 1

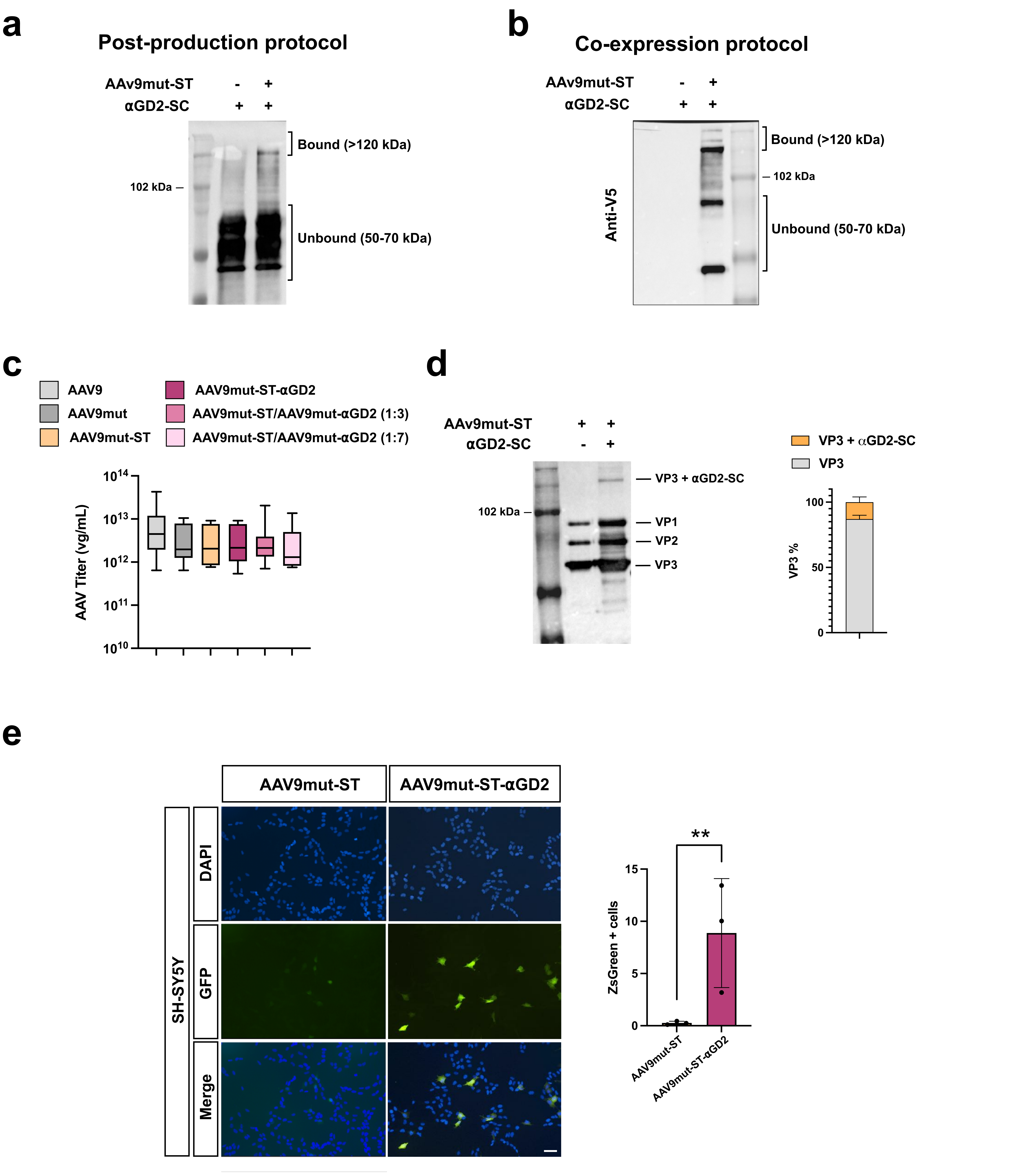

#### Supplementary Figure 2

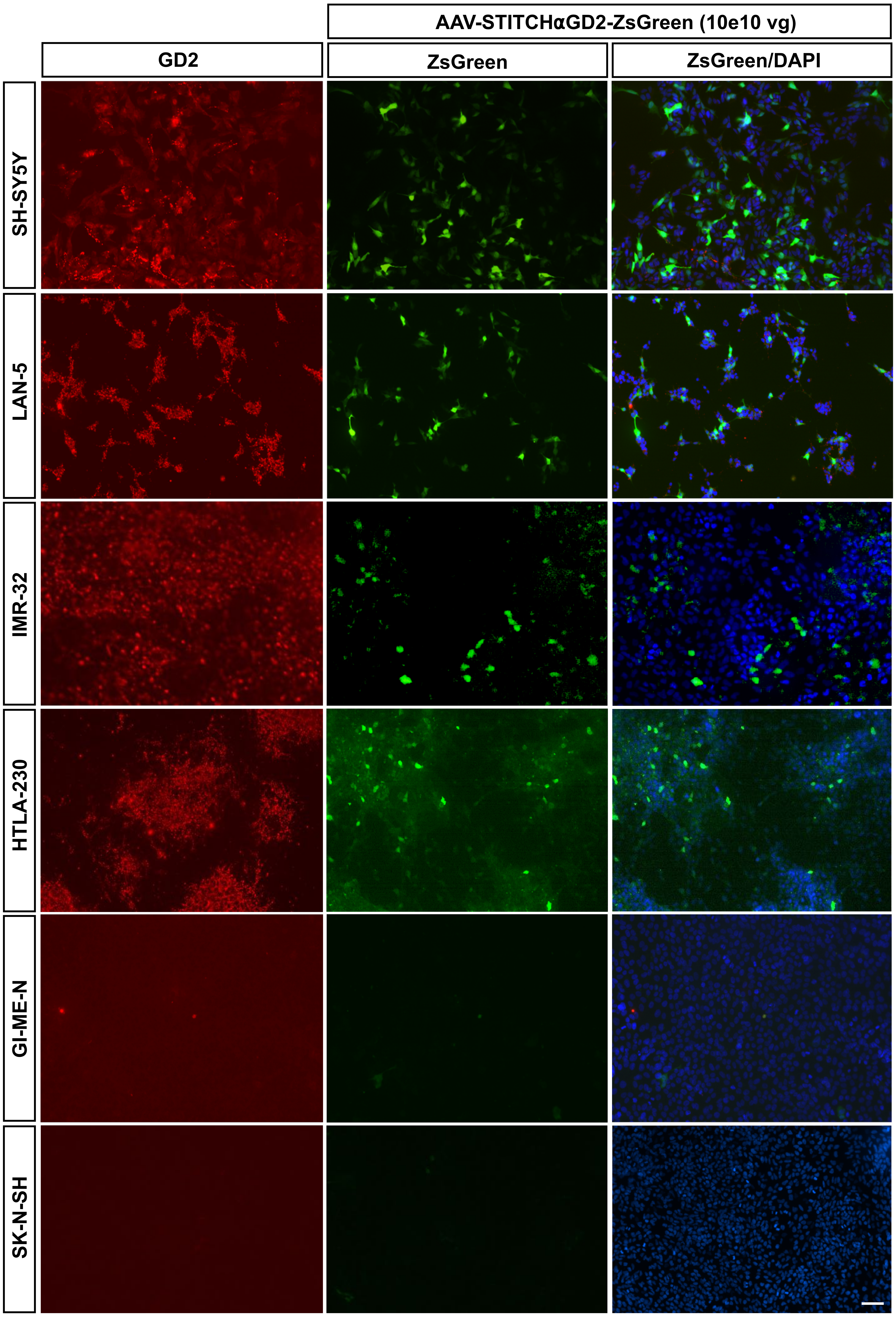


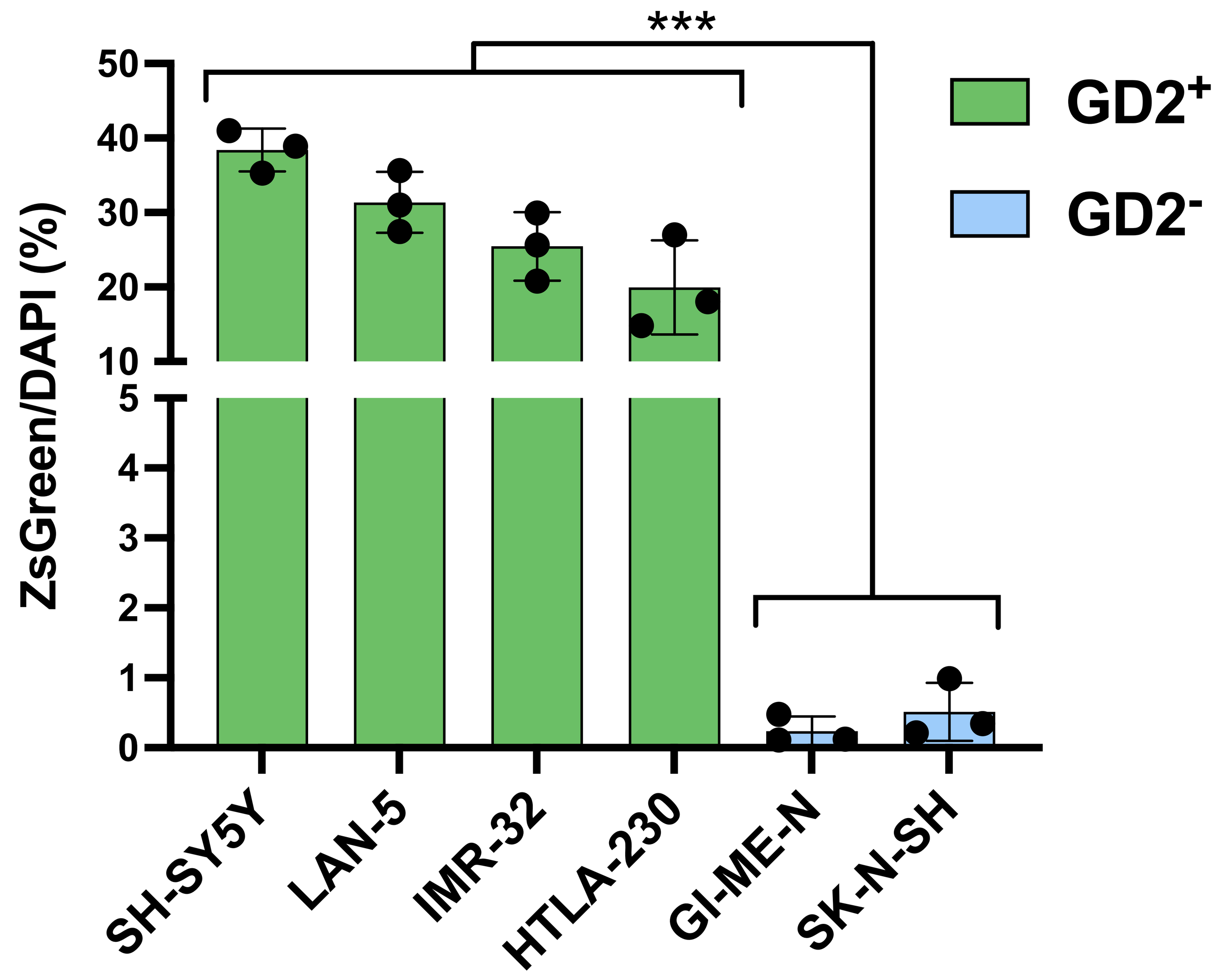

#### Supplementary Figure 3

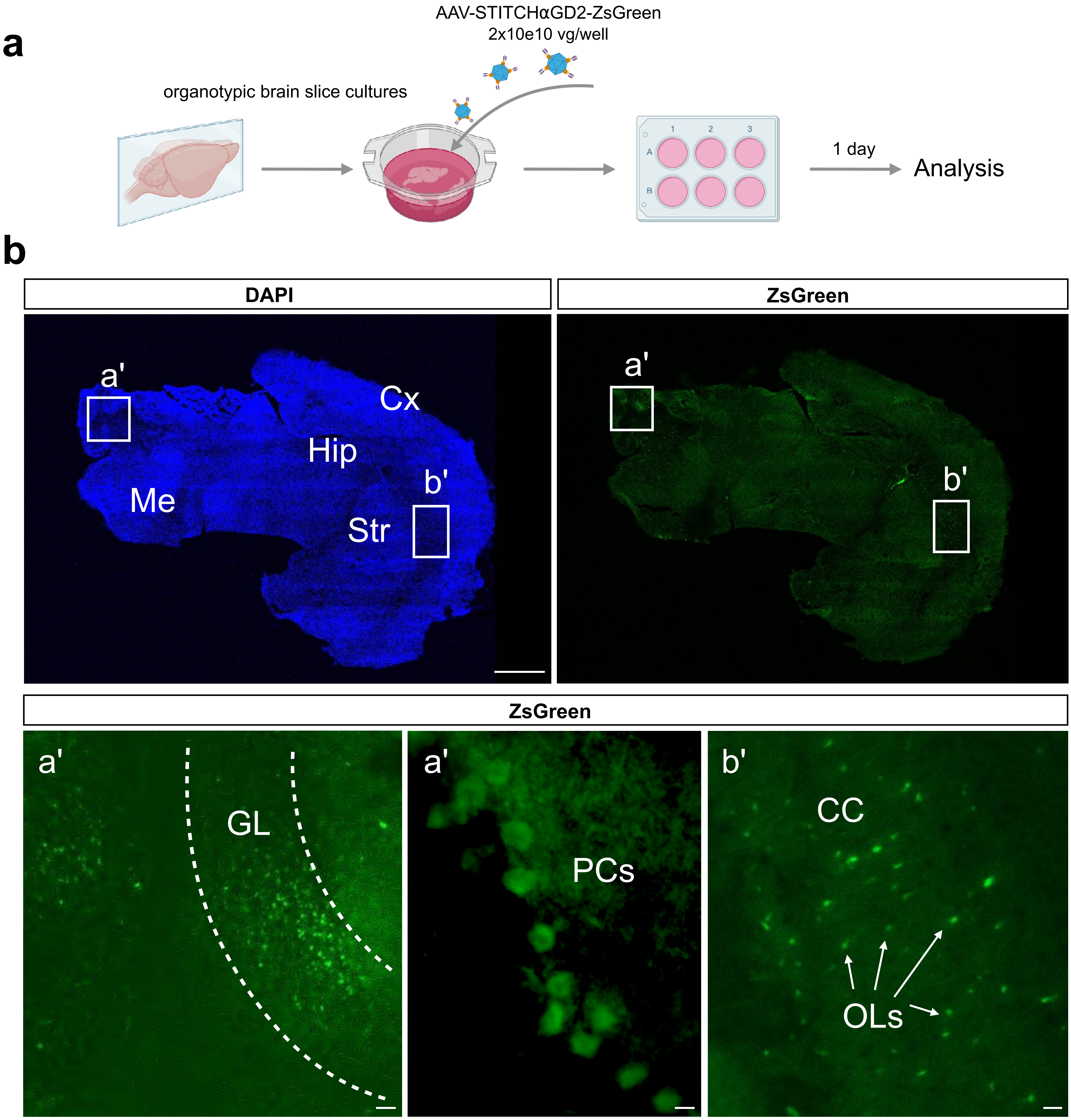

##### Supplementary Figure 4

**a**

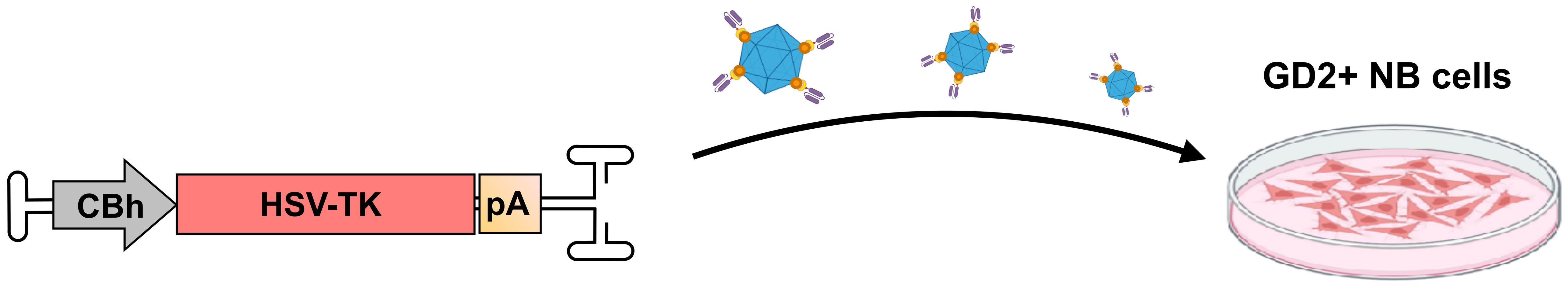


**b**

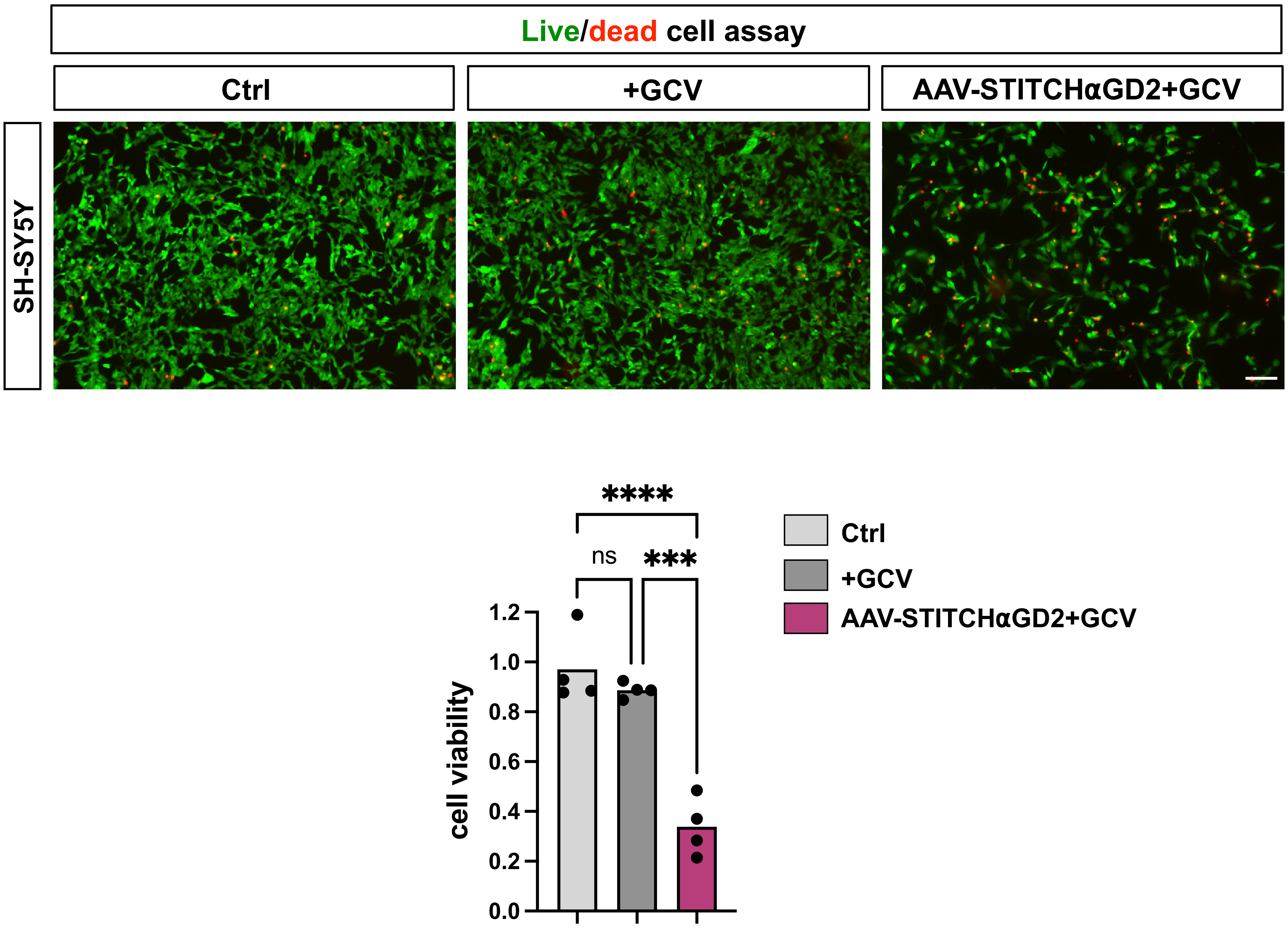
